# A shared genetic regulator of metabolism and addiction-related behavior in mice and humans

**DOI:** 10.64898/2025.12.23.696269

**Authors:** Jason A. Bubier, Michael C. Saul, Heidi S. Fisher, Price E. Dickson, Sarah A. Schoenrock, Robyn L. Ball, Anne Czechanski, Whitney Martin, Udita Datta, Leona Gagnon, Hao He, Jordyn J. VanPortfliet, Tyler Roy, Troy D. Wilcox, Laura G. Reinholdt, J. David Jentsch, A. Phillip West, Christopher L. Baker, Vivek M. Philip, Lisa M. Tarantino, Elissa J. Chesler

## Abstract

Substance use disorders and other mental health conditions often co-occur with metabolic disorders, suggesting shared biological underpinnings^1^. These heightened behavioral and physiological responses may have evolved to promote survival during resource scarcity but can become maladaptive in modern environments^2^. The genetic mechanisms linking these traits have remained elusive. Here, we show that a variable gene enhancer in mice jointly regulates genes encoding an epigenetic factor (*Eed*) and a mitochondrial enzyme (*Me3*) thereby influencing propensity to addiction-related behaviors and mitochondrial function. We further identify variation in a corresponding enhancer in humans regulating *EED* and *ME3* associated with substance use, psychiatric and metabolic disorders. These findings reveal a convergent genetic regulatory network linking mitochondrial biology to behavioral and metabolic risk, offering insight into how genetic variation in beneficial regulatory pathways can predispose individuals to substance use disorders and related conditions.

## Introduction

Substance use disorders (SUDs), including cocaine use disorder (CUD), are complex psychiatric conditions shaped by genetic and environmental factors^3^. Although many individuals encounter cocaine, only some develop compulsive use^4,5^, raising critical questions about what biological mechanisms confer vulnerability. Human studies find relations among traits such as impulsivity, novelty seeking, and risk-taking to SUDs, including CUD^6–17^, and these behaviors often precede drug exposure, suggesting a genetic basis to susceptiblity^18^. Yet, causal relationships remain difficult to establish in clinical populations due to confounding factors including polysubstance use and varied environmental exposure histories^4,19^.

Genetic variation alters susceptibility to SUD by influencing multiple behavioral and neurobiological domains, including dopamine circuit activity, synaptic plasticity, and transcriptional profiles^20^, but the underlying mechanisms remain poorly understood. Post-mortem human studies provide some insight into biological pathways, but they obscure causal effects, as they often reflect cumulative drug exposures across years, highlighting the need for experimental systems. In contrast, rodent studies offer controlled genetic and environmental conditions, enabling controlled analysis of the biology underlying addiction-like behaviors. The historical reliance on individual inbred strains, however, restricts their translational value given the variation in addiction vulnerability inhuman populations. To overcome this, we leveraged the Diversity Outbred (DO) mouse population^21^, which captures extensive genetic variation across a full range of biological pathways. These mice exhibit robust variation in exploratory and novelty-seeking behaviors that correlate with cocaine self-administration^22^, enabling genetic mapping of polygenic traits and the mechanistic dissection of addiction-related vulnerability^23^.

Here we sought to identify the genetic factors mediating the relationship between traits associated with behavioral vulnerability to CUD and relevant phenotypes in mice. To this end, we applied a multi-dimensional genetic mapping analysis of mouse behavior. The convergence of behavioral, genetic, and molecular data revealed a candidate mechanism associated with mitochondrial function that we evaluated experimentally and extrapolated to behavioral and metabolic characteristics in mice and human populations using bioinformatics. Together, these findings reveal a shared genetic mechanism connecting energy metabolism and behavioral susceptibility, offering insights into how advantageous pathways can predispose individuals to SUD, including cocaine addiction.

## Methods

See Supplementary Extended Methods for full description.

### Animals

Diversity Outbred (DO) mice were obtained from The Jackson Laboratory and characterized across multiple behavioral and molecular assays. All procedures were approved by the Institutional Animal Care and Use Committee.

### Reference trait phenotyping study design

To address carry-over and other challenges inherent to longitudinal study designs, we developed a canonical correlation-based framework to project behavioral and drug-response traits across independently assayed cohorts^24^. This approach enabled the integration of drug naïve, exploratory behavior phenotypes measured across the full population with exposure-related phenotypes, such as cocaine self-administration and behavioral sensitization, measured on subsets of the population.

Mice were tested in a battery of behavioral assays, including open field, light/dark box, hole board, and novel place preference to assess novelty seeking, risk-taking, and exploratory behavior. These “reference” traits were used to construct predictive indices for treatment response “target” traits that were used in downstream analyses. A subset of mice underwent multi-phase cocaine intravenous self-administration (IVSA) testing, including acquisition, dose-response, extinction, and cued reinstatement phases. Another subset of mice was tested for locomotor sensitization to repeated cocaine exposure (COC), alongside saline-treated controls (SAL). Behavioral metrics were derived from activity patterns across a 19-day protocol.

Reference trait analysis was used to impute addiction-related traits across the full DO cohort using canonical correlation methods.

### Genotyping, genetic mapping, and candidate gene prioritization

Genomic DNA was genotyped using the GigaMUGA array. Quantitative trait loci (QTL) mapping was performed using R/qtl2 to identify loci associated with imputed behavioral traits. Pleiotropy was assessed across traits.

Genes within QTL intervals were ranked based on cis-eQTL effects, LOD score correlation, and founder allele effect concordance.

### Single-nucleus multiomic profiling

snRNA-seq and ATAC-seq were performed on striatal tissue from DO founder strains to identify cell-type specific regulatory elements.

### CRISPR editing, RNA sequencing, and mitochondrial imaging

Putative enhancer regions were deleted in mESCs from C57BL/6J and NOD/ShiLtJ strains using CRISPR/Cas9. Edited lines were differentiated into neural progenitor cells (NPCs) for transcriptomic and imaging analyses. RNA sequencing was performed on NPCs with and without the deletion to assess expression of *Eed*, *Me3*, and neuronal markers. Simliarly, immunofluorescence microscopy was used to assess mitochondrial morphology and protein expression in NPCs.

### Bioinformatics

Cross-species and human regulatory analyses were performed using GeneWeaver, GeneHancer, RegulomeDB, GWAS Catalog, and Mouse Phenome databases to identify convergenet mechanisms.

## Results

### Reference trait analysis indicates a shared genetic basis for multiple addiction related behaviors

Canonical correlation analysis within each testing cohort revealed relations among baseline behaviors and IVSA, COC and SAL. The first canonical correlation for each target trait was largely correlated with sex differences in the behaviors (IVSA |ρ| = 0.09, COC |ρ| = 0.12, SAL |ρ| = 0.13; Table S3). The second canonical correlation reflected more well-distributed (IVSA |ρ| range = 0.008-0.06, COC |ρ| range = 0.006-0.06, SAL |ρ| range = 0.004-0.08) contributions of multiple reference and target traits and thus was used for deriving index scores (Table S3) for the target traits, cocaine IVSA, COC and SAL for the entire population of DO mice, including those mice that had not been tested for self-administration or behavioral sensitization (Figure 1a). In three separate canonical correlation analyses, using the second canonical correlate in each, we found that both distinct and overlapping novelty response and exploratory behavior variables were predictive of IVSA (Figure 1b), COC (Figure 1c) and SAL traits (Figure 1d).

**Figure 1.**
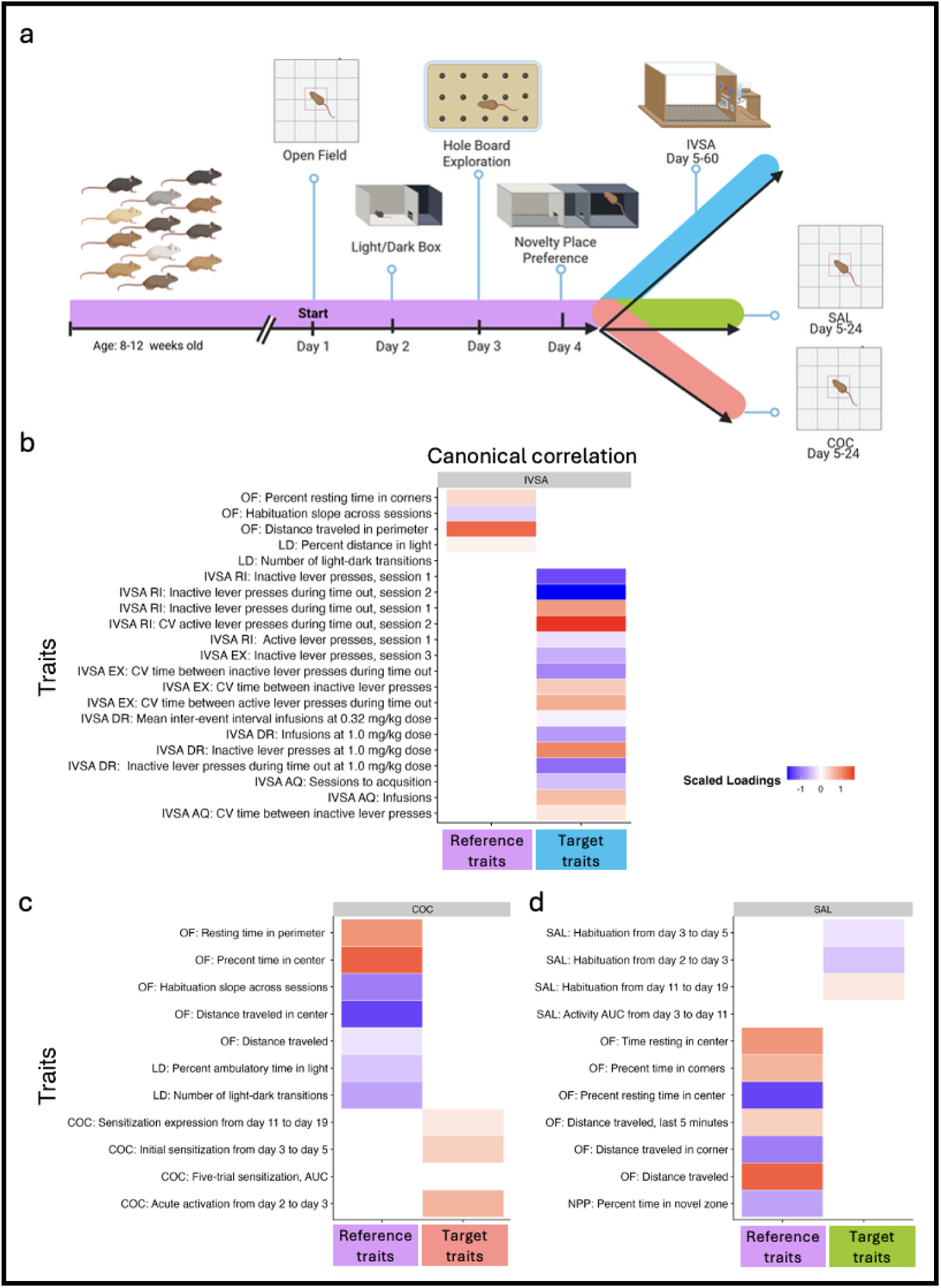
Characterization and correlation of predisposing and addiction-related behaviors in a diverse mouse population. (a) Diversity Outbred mice were first phenotyped for reference traits (open field, light-dark box, hole board, novel place preference), then subset into groups to measure target traits: cocaine intravenous self-administration (IVSA), cocaine sensitization (COC), and saline habituation (SAL). Canonical correlations for the target traits (b) IVSA and (c) COC and (d) SAL and associated correlation weights indicated by scaled loadings, blue represents negative association and red positive. Canonical correlations among each of the three target traits show shared regulation of phenotypic variation despite being composed from different combinations of loadings.

### Genetic mapping of IVSA, COC and SAL reveal shared regulatory loci

We detected significant QTL for each the three derived target traits (Table 1) on Chromosome 7: IVSA (*Civsa1* - cocaine intravenous self-administration; Figure 2a, top panel), COC (*Cocia22* cocaine-induced activity, 22; Figure 2a, middle panel), and SAL (Figure 2a, bottom panel).

**Figure 2.**
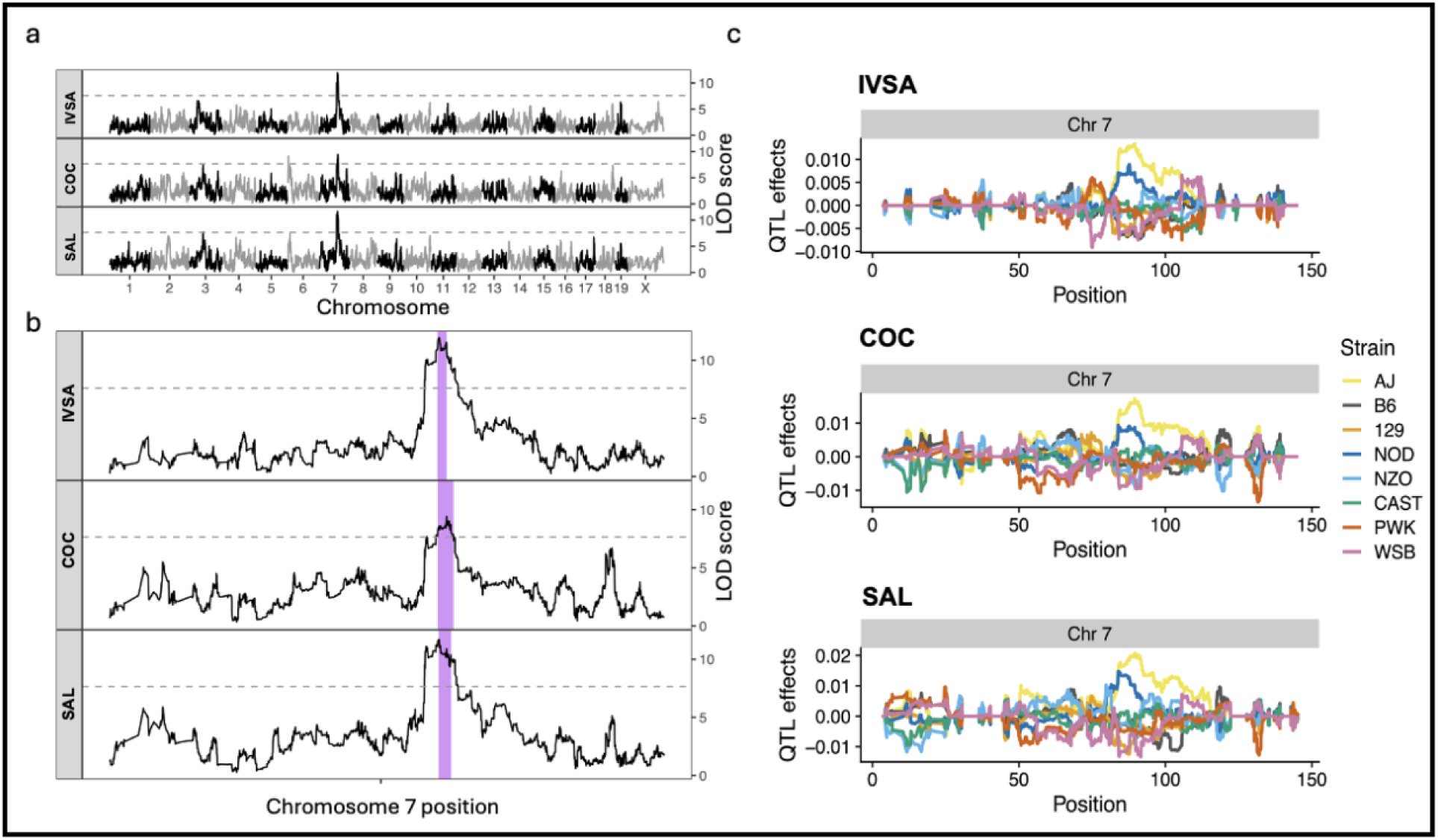
Genetic mapping of cocaine intravenous self-administration (IVSA), cocaine sensitization (COC), and saline habituation (SAL). Quantitative trait loci (QTL) mapping of reference traits across (a) the genome and (b) chromosome 7, with confidence intervals depicted as purple bars surrounding peak makers and suggesting shared genetic regulation of the three traits. (c) QTL allele effects across chromosome 7 for the reference traits showing an increase of effect of A/J (yellow line) and NOD/ShiLtJ (NOD, dark blue lines) alleles and decrease of effects of other alleles including C57BL/6J (B6, dark grey bars), 129S1/SvImJ (129, orange), NZO (NZO/HILtJ, light blue), CAS (CAST/EiJ, green), PWK (PWK/PhJ, red), and WSB (WSB/EiJ, lavender).

**Table 1.** QTL mapping peaks with genome-wide permutation-based signficance < 0.05.

| Phenotype | LOD score | Chromosome | Peak Position (GRCm38) | 95% CI Low (GRCm38) | 95% CI High (GRCm38) |
| --- | --- | --- | --- | --- | --- |
| IVSA | 13.25 | 7 | 87650445 | 87127600 | 89522474 |
| SAL | 11.67 | 7 | 87464127 | 84039573 | 90658691 |
| IVSA | 9.57 | 10 | 126992303 | 126910230 | 128357832 |
| COC | 9.41 | 7 | 89522474 | 87244659 | 91284113 |
| COC | 9.12 | 6 | 5211712 | 4997746 | 5628980 |
| SAL | 7.74 | 3 | 69260258 | 66826817 | 76848854 |

Pairwise pleiotropy analyses of these three derived traits (Figure S1) indicated that they are regulated by the same genetic factors. This consensus QTL (*Cocrb17*- cocaine related behavior 17) on Chromosome 7 spans a region from 84,039,573 bp to 91,284,113 bp (GRCm38 assembly; Figure 2b) based on overlapping confidence intervals of the IVSA, COC, and SAL QTLs. C57BL/6J and NOD/ShiLtJ allelic effects are divergent for all three phenotypes (Figure 2c). Linkage disequilibrium analysis using mutual information coefficient as a measure of entropy shows strong linkage between the COC peak marker at Chr7:87,464,127 and the SAL marker at Chr7:89,522,474, encompassing the IVSA marker at Chr7:87,650,445 (Figure S2). Additional QTLs were identified on Chromosome 10 for IVSA, Chromosome 6 for COC, and Chromosome 3 for SAL (Table 1).

### Integrative analyses implicate an expression regulatory network involving multiple genes

The consensus QTL interval spanning 7.25 Mb contains 41 protein-coding genes. We prioritized positional candidate genes based on evidence for a significant cis-eQTL at a threshold of FDR < 0.05, using previously published bulk striatal RNASeq data^25^ to focus on 15 genes, of which, only four (*Eed*, *Me3*, *Fah* and *Ccdc81*) had a cis-eQTL with an allelic segregation pattern that matched the QTL allelic effect segregation pattern (divergent C57BL/6J and NOD/ShiLtJ effects; Figure S3).

### Single cell multiomics identifies a common regulator of Eed and Me3

Single cell RNAseq and ATACseq analyses of striatal tissue from the 8 DO founder strains^26^ revealed 18 previously characterized striatal cell types (Figure 3a; N = 55,672 total nuclei). The top five clusters include medium spiny neurons (MSNs) expressing either *Drd1* (N = 15,395) or *Drd2* (N = 13,203) dopamine receptors, oligodendrocytes (N = 12,066), astrocytes (N = 4,748), and microglia (N = 2,638). Both *Me3* and *Eed* were expressed in all five cell types. *Me3* was highly expressed in both *Drd1* and *Drd2* cell types (normalized expression > 0.50; Figure 3b) compared with oligodendrocytes, astrocytes and microglia (normalized expression < 0.25; Figure 3b). Within the consensus QTL (*Cocrb17*), several open chromatin regions were identified that showed increased accessibility in *Drd1* and *Drd2* medium spiny neurons, compared to oligodendrocytes, astrocytes, and microglia (Figure 3c). Focusing on *Drd1* MSNs, we identified significant putative co-variation (FDR < 0.01) for two open chromatin regions within a 3’ intron of the *Me3* coding region with both *Me3* and *Eed*, suggesting their function as enhancers. Interactions between *Me3* and *Prss23,* and *Me3* and *Ccdc81* were also identified in *Drd2* neurons. *Me3*, *Eed, Ccdc81* share predicted cis-regulatory elements that are significantly associated (FDR < 0.05) in *Drd2* medium spiny neurons (Figure 3d). Significant interactions between *Me3* and *Fzd4,* and *Fzd4* and *Prss23* were also identified in *Drd1* MSNs; however, neither *Fzd4* nor *Prss23* were highly ranked candidate genes based on the criteria described above. Moreover, *Fah* was not identified in either *Drd1* or *Drd2* MSNs and therefore narrowed our focus from four focal genes (*Eed*, *Me3*, *Fah* and *Ccdc81*) to *Me3, Eed,* and *Ccdc81.* Finally, we identified two genetic variants within the QTL containing *Me3, Eed,* and *Ccdc81* and their eQTLs: *Rr607* (‘Peak A’; Chr7:89,837,035 – 89,837,873 bp), located 674 bp from the second of the two open chromatin regions and Peak B (Chr 7:89,841,645 - 89,842,644 bp).

**Figure 3.**
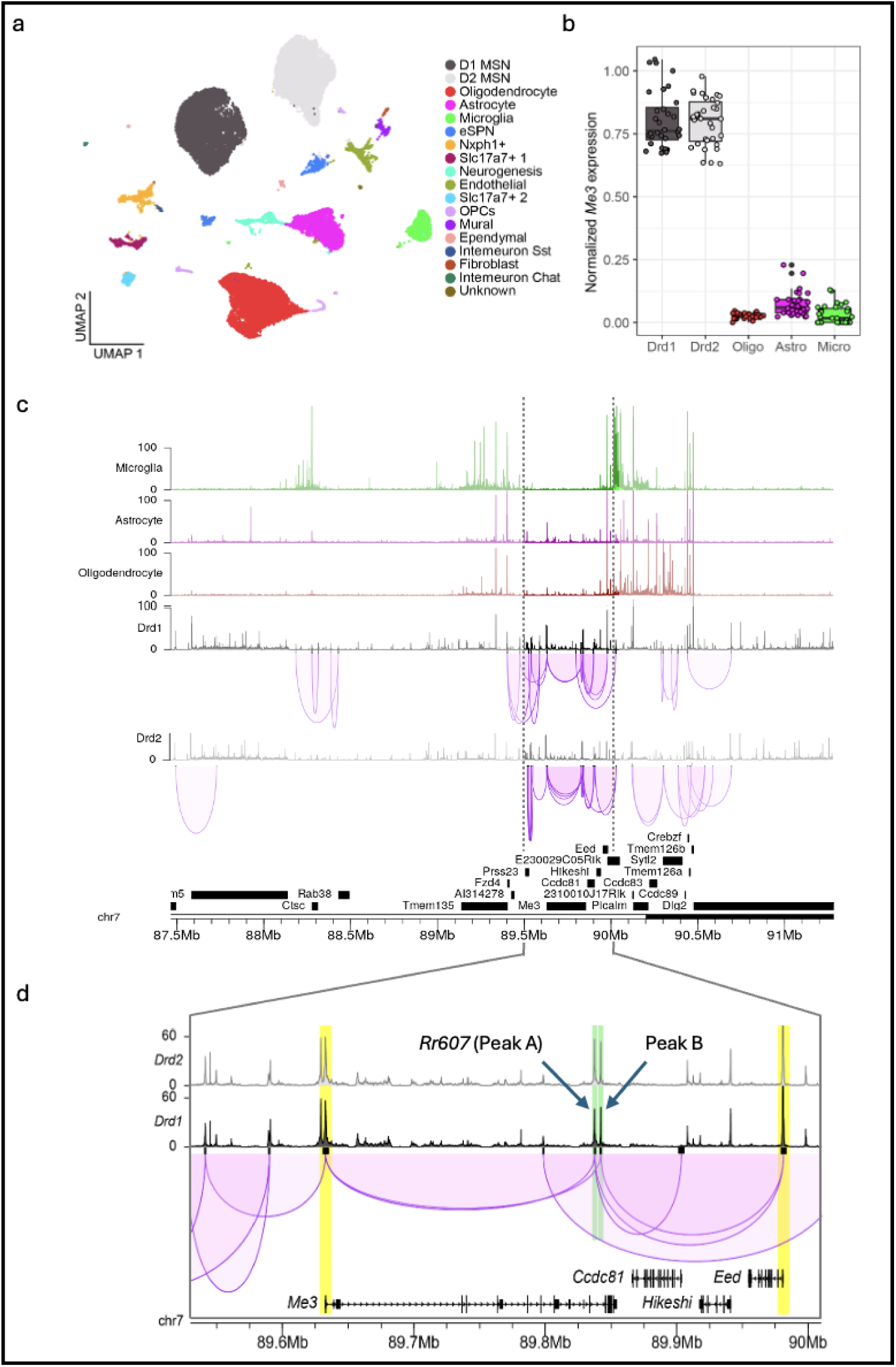
Single-nuclear multi-omic profiling of the founder striatum. (a) UMAP of 55,672 nuclei reveals predominant cell-types based on expression patterns with notable clusters from *Drd1* medium spiny neurons (D1 MSN, black) and *Drd1* (D2 MSN, grey). (b) Normalized *Me3* expression is highly expressed in *Drd1* (black) and *Drd2* (grey) MSN compared to other cell types. (c) Enhancer-promoter interactions in *Drd1* and *Drd2* MSN within (a) shared QTL interval of IVSA, COC, and SAL, and (b) focal region, in which *Me3* and *Eed* (yellow shading) share predicted cis-regulatory elements as indicated by the purple looping patterns. *Rr607* (Peak A) and Peak B, both shaded in green, are predicted enhancers that of *Eed* and *Me3; Rr607* is also linked to *Ccdc81*. Both peaks are unique to *Drd1* and *Drd2* cell populations and absent in cell populations from microglia, astrocytes and oligodendrocytes.

Strain specific (C57BL/6J and NOD/ShiLtJ) multi-omic data revealed high and low expression based on the eQTL allele effect and behavioral QTL allele effects (Figures S1A-C)^27^. We observed higher expression of *Me3* in *Drd1* (*p_adj_* < 0.001; Figure 4a) and *Drd2* (*p_adj_* = 0.035; Figure 4b) expressing MSNs in C57BL/6J relative to NOD/ShiLtJ.ShiLtJ. In contrast, we observed higher expression of *Eed* in *Drd1* (*p_adj_* < 0.001, Figure 4a) but not in *Drd2* (*p_adj_* > 0.05; Figure 4b) neurons, and higher expression of *Ccdc81* (*Drd1*: *p_adj_*< 0.001; *Drd2*: *p_adj_* = 0.035; Figure 4), in NOD/ShiLtJ relative to C57BL/6J in *Drd1* and *Drd2* expressing MSNs. We found a significant strain specific interaction at *Rr607* (Peak A; *Drd1* expressing MSN, *p_adj_* = 0.005; *Drd2* MSNs: *p_adj_*= 0.016; Figure 4), with NOD/ShiLtJ exhibiting high expression, whereas in Peak B we see no strain differences (*Drd1* MSNs: *p_adj_* > 0.05; *Drd2* MSNs: *p_adj_* > 0.05; Figure 4) and therefore excluded Peak B as a putative causal region. Together, single-nuclei multi-omic analysis, combined with predicted regulatory links, implicates a single putative enhancer, *Rr607*, as the cis-regulatory element driving *Me3* and *Eed* expression, along with other genes in the region and cocaine-related behaviors.

**Figure 4.**
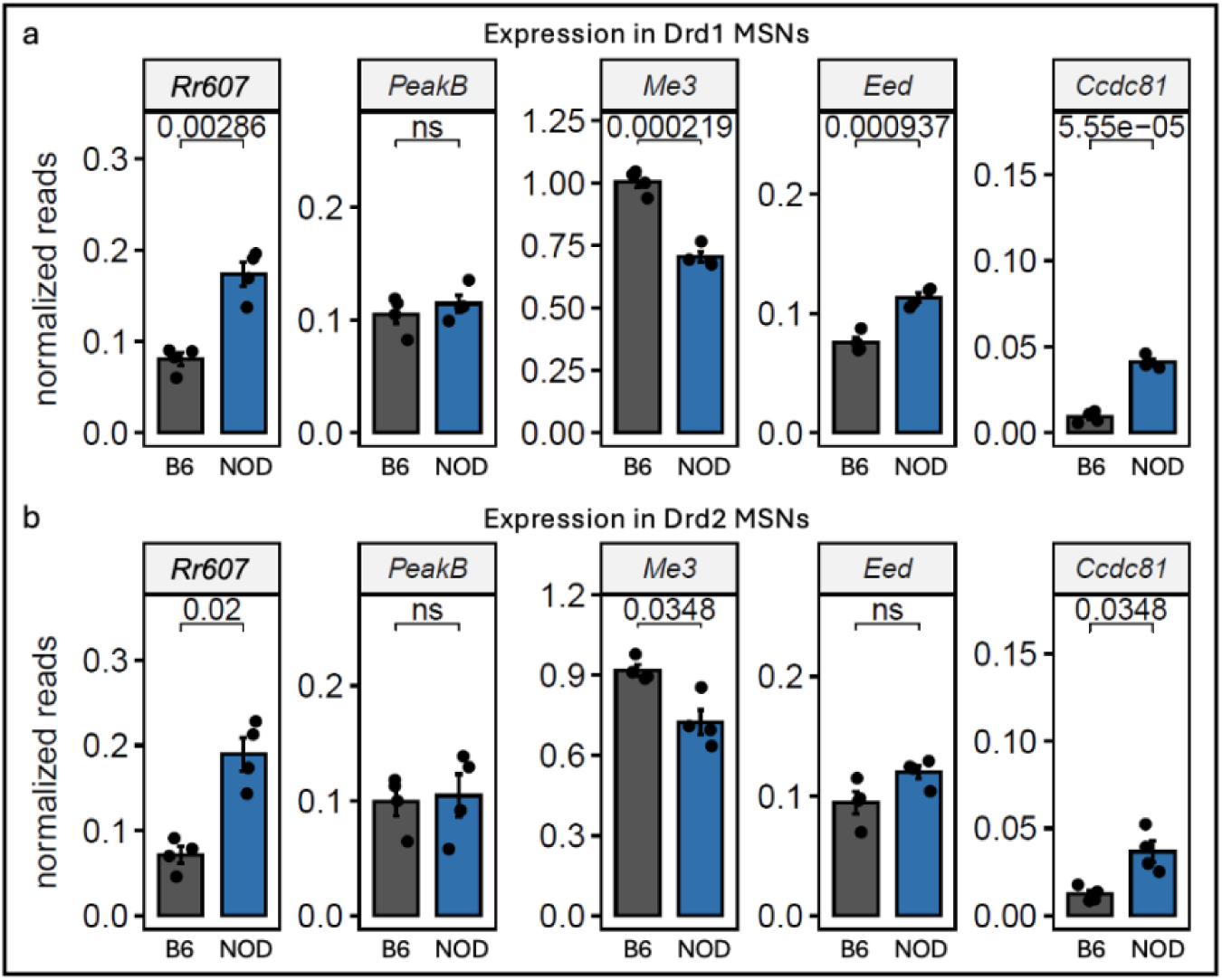
Strain-specific regulatory peak and gene expression in (a) *Drd1* and (b) *Drd2* medium spiny neurons (MSNs) across C57BL/6J (B6, dark grey bars) and NOD/ShiLtJ (NOD, dark blue bars) strains. ATAC-seq reads indicate that *Rr607* (Peak A) has a greater number of relative contacts with the predicted enhancer regions in NOD/ShiLtJ compared to C57BL/6J in both *Drd1* and *Drd2* MSN, whereas Peak B does not differ. RNA-seq reads suggest significant gene expression differences in *Eed, Me3* and *Ccdc81* in *Drd1* MSN and less profound effects in *Drd2* MSN.

### C57BL/6J and NOD/ShiLtJ show differential expression of Eed and Me3 modulated by Rr607

When we compared strain differences in wild-type neural precursor cells (NPCs), we observed that NOD/ShiLtJ cells have higher expression of *Eed* (logFC = 0.29, *p_adj_* = 6.6 x 10^-5^) and *Me3* (logFC = 2.50, *p_adj_* = 3.3 x 10^-33^), but no difference in expression of *Ccdc81* (logFC = -0.09, *p_adj_* = 0.923), compared to C57BL/6J cells (Figure 5a). The genetic ablation of *Rr607* had no effect on *Eed* (logFC = 0.01, *p_adj_* = 0.950), *Me3* (logFC = 0.39, *p_adj_* = 0.097), or *Ccdc81* (logFC = 0.62, *p_adj_* = 0.61) expression in C57BL/6J cells (Figure 5a). However, in NOD/ShiLtJ cells, *Rr607* deletion resulted in a significant decrease in *Me3* expression (logFC = -2.53, *p_adj_*< 0.00001; Figure 5a), reducing it to a level similar to C57BL/6J wild-type expression. The deletion of *Rr607* in NOD/ShiLtJ cells also reduced the expression of *Eed* (logFC = -0.16, *p_adj_* = 0.005; Figure 5a) but had no effect on *Ccdc81* expression (logFC = 0.31, *p_adj_* = 0.76; Figure 5a). For both *Me3* and *Eed*, deletion of *Rr607* resulted in expression levels that were not significantly different than those observed in wild-type C57BL/6J cells (*Me3:* logFC = 0.39, *p_adj_* = 0.22; *Eed*: logFC = -0.01, *p_adj_* = 0.95; Figure 5a). These findings further narrowed our focus from three focal genes (*Eed*, *Me3* and *Ccdc81*) to *Me3* and *Eed*.

**Figure 5.**
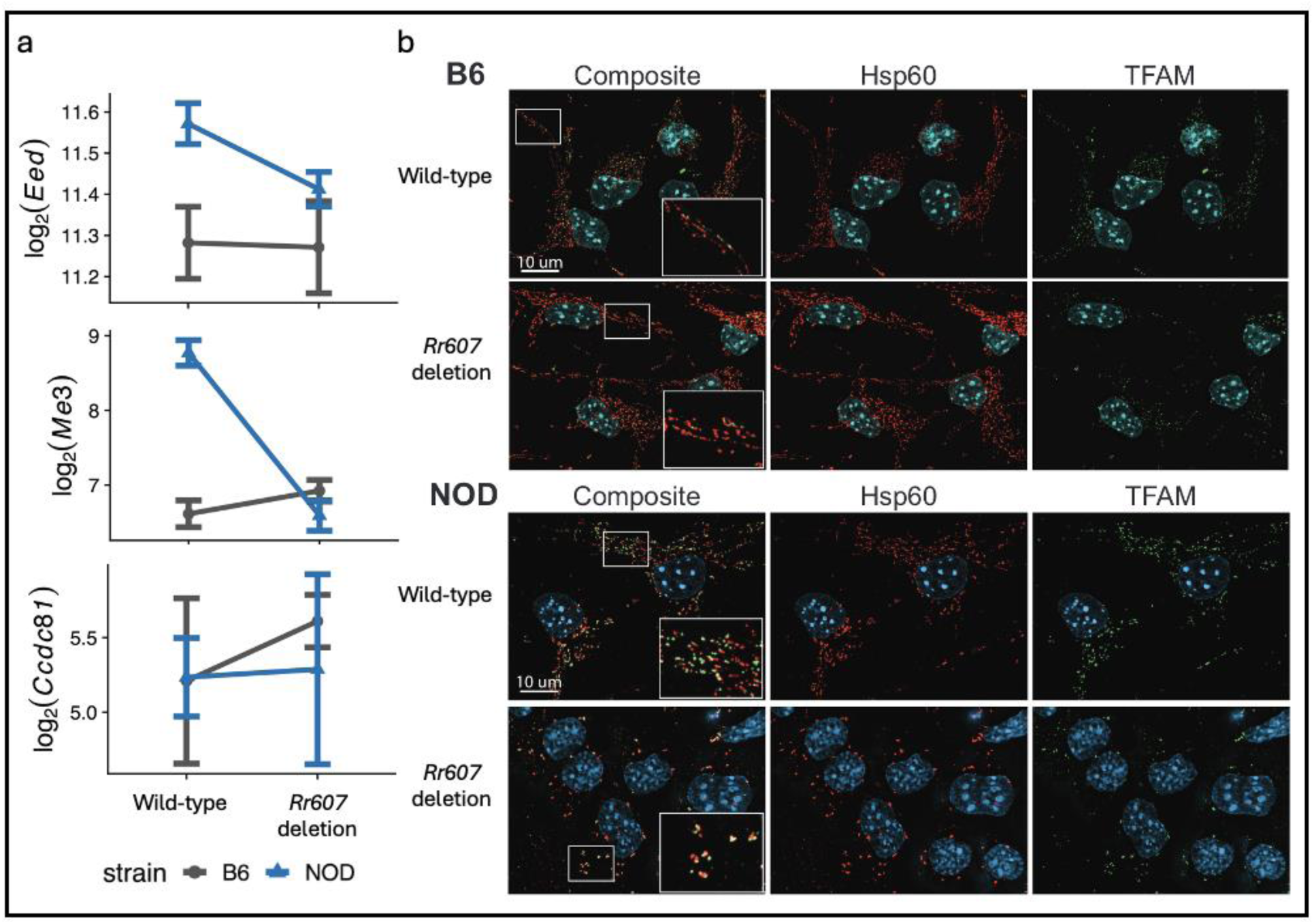
Strain-specific effects of *Rr607* enhancer deletion in C57BL/6J (B6, dark blue) and NOD/ShiLtJ (NOD; dark grey) strains. (a) *Me3, Eed,* and *Ccdc81* expression in response to *Rr607* deletion in neuronal progenitor cells (NPCs) derived from the striatum of C57BL/6J and NOD/ShiLtJ mice. Strain differences in *Eed* and *Me3* expression are eliminated by the deletion of the enhancer, with no effect on *Ccdc81*. (b) Representative immunofluorescence images of NPCs generated from C57BL/6J and NOD/ShiLtJ backgrounds with wild-type and *Rr607* enhancer deletion. Cells are stained with DAPI to indicate the nucleus, heat shock protein 60 (Hsp60, red) to mark the mitochondrial networks, and mitochondrial transcription factor A (TFAM, green) to mark mitochondrial nucleoids. *Rr607* enhancer deletion in NOD/ShiLtJ, but not C57BL/6J, are suggestive of a mitochondrial fission.

### NOD/ShiLtJ NPCs exhibit fragmented mitochondrial networks and loss of TFAM after Rr607 deletion

*Me3* and *Eed* are genes that interact to regulate cellular metabolism and mitochondrial processes, some of which are associated with cocaine response^28^. *Me3* encodes for a mitochondrial-specific isoform of malic enzyme that catalyzes oxidative decarboxylation of malate to pyruvate with an NADP(+) cofactor and has a known role in regulating mitochondrial function and oxidative stress^29^. *Eed* encodes a polycomb repressive complex PRC2 protein that binds to H3K27me3^30^ and is involved in transcriptional repression, which influences reactive oxygen species (ROS) production via regulation of *Me3* and mitochondrial activity^31^. ROS production is essential for synaptic plasticity^32^ and the necessary cytoskeletal restructuring that occurs in cocaine-related responses in rodents^33^. A recent report linked malic enzyme expression to mitochondrial DNA replication and mitochondrial translation^34^. Therefore, to assess effects of the enhancer on mitochondria function, we employed immunofluorescence microscopy to assess expression of the mitochondrial matrix protein, Transcription Factor A, Mitochondrial (TFAM; labeled in green, Figure 5b), which is an essential regulator of mitochondrial DNA replication, transcription and packaging^35^. We found that the *Rr607* (Peak A) enhancer deletion altered relative TFAM expression (log of TFAM area normalized to cell area) in a strain-specific manner in NOD/ShiLtJ and C57BL/6J NPCs (*F*_1, 42.09_ = 19.31, *p* < 0.0001, Figure S5a), such that NOD/ShiLtJ NPCs showed a dramatic decrease in TFAM with the deletion compared to wild-type (*t* = 5.58, *df* = 42, *p* < 0.0001, Figure S5b), yet C57BL/6J NPCs did not differ between wild-type and *Rr607* deletion *(t* = -0.63, *df* = 42, *p* = 0.53, Figure S5b).

Thus, *Rr607* deletion results in a phenotype consistent with altered mitochondrial DNA abundance and function in the NOD/ShiLtJ background, which can lead to altered cellular metabolism.

To assess mitochondrial networks, we used an antibody against the mitochondrial heat shock protein 60 (HSP60; labeled in red, Figure 5b). We found that the effect of the *Rr607* deletion altered relative mitochondrial count (log of count normalized to cell area) in a stain-specific manner (*F*_1, 42.00_ = -2.10, *p* = 0.042, Figure S5c); specifically, C57BL/6J NPCs with the *Rr607* deletion have relatively more mitochondria than wild-type (*t* = -2.03, *df* = 42, *p* = 0.049, Figure S5d), and NOD/ShiLtJ NPCs do not differ (*t* = 0.94, *df* = 42, *p* = 0.35, Figure S5d). Moreover, the *Rr607* deletion altered the relative area of mitochondrial networks (log of area normalized to cell area) in a stain-specific manner (*F*_1, 42.00_ = 10.91, *p* = 0.0019, Figure S5e). Specifically, C57BL/6J NPCs with the *Rr607* deletion have relatively larger mitochondrial networks than wild-type (*t* = -2.80, *df* = 42, *p* = 0.0078, Figure S5f), and NOD/ShiLtJ NPCs with the *Rr607* deletion have slightly smaller mitochondrial networks than wild-type (*t* = 1.87, *df* = 42, *p* = 0.069, Figure S5f). Moreover, we also found wild-type NOD/ShiLtJ NPCs exhibit significantly larger mitochondrial networks than wild-type C57BL/6J NPCs (*t* = -2.437, *df* = 42, *p* = 0.019, Figure S5f).

### Identification of additional mouse phenotypes associated with Rr607

We calculated Lewontin’s D’ measure of linkage disequilibrium (LD) between the six variants within *Rr607* and 71 nearby variants typed in the two focal strains, C57BL/6J and NOD/ShiLtJ, for which data exists in Mouse Phenome Database (D’ range: 0.027 - 1; median = 0.941). Of these, we filtered for the 67 variants in LD with at least one of the six variants (D’ ≥ 0.80), and further filtered for the 53 variants that also have different allelic states between C57BL/6J and NOD/ShiLtJ. These 53 variants were associated with 304 phenotypic traits in the Mouse Phenome Database, which included multiple behavioral and metabolic traits such as alcohol intake, response to nicotine, anxiety-like behavior, abnormal sleep patterns, body size, adipose tissue and glucose tolerance (Mouse PheWAS: Figure 6, Table S4).

**Figure 6.**
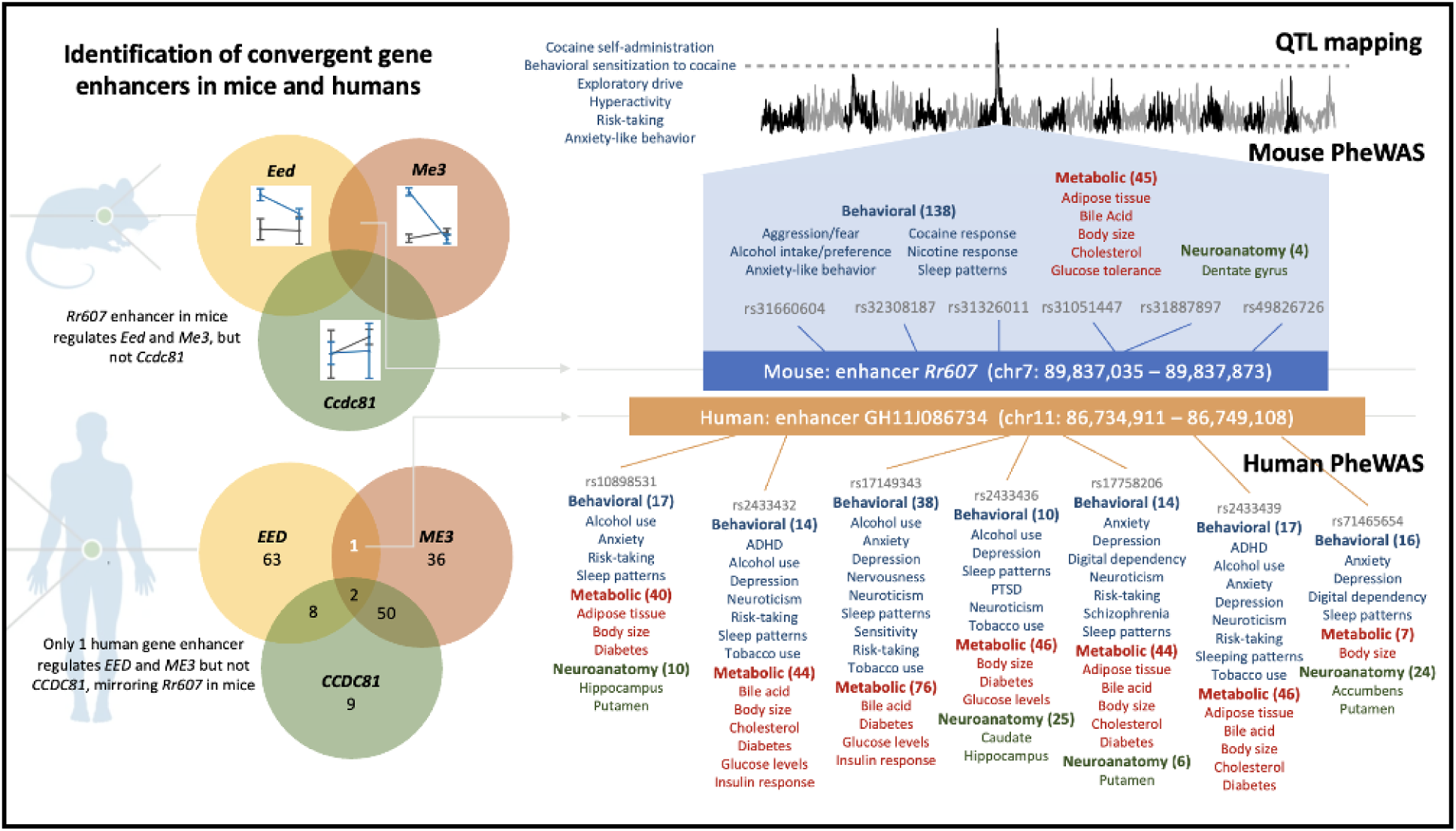
Enhancer variation in mice and humans is linked to similar patterns of behavioral (blue), metabolic (red), and neuroanatomy (green) phenotypes. Genetic mapping in Diversity Outbred mice identified a QTL that jointly regulates cocaine self-administration, behavioral sensitization to cocaine, and other predisposing behavior traits. A Mouse Phenome Database search of variants linked to a causal enhancer within the QTL revealed a wide range of behavioral patterns related drug response, anxiety-like behavior and sleep patterns, metabolic traits including glucose tolerance, cholesterol levels, and fat storage. A convergent enhancer in humans that similarly regulates expression of *ME3* and *EED*, but not *CCDC81,* and shows association with a range of addiction-related behaviors and other psychiatric traits including anxiety, depression, abnormal sleep patterns and risk-taking, as well as metabolic traits suchsuch as body size and diabetes, and associated neuroanatomy phenotypes.

### Identification of a conserved joint regulatory locus in humans

Queries of GeneWeaver revealed two different human studies showing differential regulation of both EED and ME3 in people with CUD vs control groups. First, in RNA-seq of dorsolateral prefrontal cortex, *ME3* (logFC = -4.0307) and *EED* (logFC = -26.774) were differentially expressed; however, neither were within with any of the weighted gene co-expression network analysis modules in that analysis^36^. Second, in a single cell RNA-seq analysis of the caudate nucleus in humans, among the genes differentially expressed between people with CUD and control samples within *Drd1* expressing MSNs were *EED* (logFC = 0.045658; *p_adj_* = 1.09 x 10^-4^) and *ME3* (log fold change = -0.32291; *p_adj_*= 3.74 x 10^-44^). They were also differentially expressed in *Drd2* expressing MSNs (*EED:* logFC = 0.053466, *p_adj_* = 5.74 x 10^-7^; *ME3:* logFC = - 0.37058 and *p_adj_*= 2.89 x 10^-70^). Consistent with the single-cell multi-omics, neither ME3 nor EED were differentially expressed in astrocytes, microglia, oligodendrocytes, or oligodendrocyte progenitor cells^37^.

Based on this evidence for the involvement of EED and ME3 in cocaine response in humans, we sought a convergent regulatory element. We identified three enhancers, GH11J086589, GH11J086475, and GH11J086734 as joint regulators of *EED* and *ME3* in humans in the GeneHancer^38^ database, then further focused on GH11J086734 because we found that deletion of the enhancer *Rr607* (Peak A) in mouse NPCs did not alter expression of *Ccdc81,* suggesting that GH11J086734 is most similiar to the mouse *Rr607* enhancer. We prioritized seven variants within GH11J086734 in the dbSNP database^39^ within RegulomeDB^40^ (rs17149343, rs17758206, rs2433439, rs10898531, rs2433436, rs2433432, rs71465654) to query existing PheWAS data in GWAS Atlas^41^. We observed PheWAS hits associated with alcohol consumption, anxiety, risk-taking behaviors, and abnormal sleep patterns, as well as neurological and metabolic traits including obesity- and diabetes-related phenotypes (Human PheWAS Figure 6; Table S5).

## Discussion

Collectively, our studies demonstrate a shared regulatory mechanism for addiction and metabolic traits in mice and humans. We identified a genetic locus in mice that jointly regulates cocaine self-administration, sensitization, and behavioral traits such as novelty-seeking, anxiety-like responses, and risk-taking. Muli-omic analyses revealed a regulatory enhancer region, *Rr607,* influences expression of two genes: *Me3*, which controls mitochondrial energy metabolism, and *Eed*, a core component of the Polycomb Repressive Complex 2 that orchestrates neuronal development. Both genes are expressed in dopamine receptor-positive (*Drd1* and *Drd2)* MSNs, the principal cell type of striatal circuits mediating reward and addiction. Single-cell ATAC-seq and RNA-seq demonstrated enhancer accessibility and transcriptional co-regulation in *Drd1*- and *Drd2*-expressing MSNs, implicating this locus in pathways directly relevant to cocaine response. Deletion of the *Rr607* enhancer altered the expression of both genes and disrupted mitochondrial networks linking mitochondrial dynamics to addiction-related behaviors^28,42^.

The *Rr607* enhancer’s pleiotropic effects span behavioral and metabolic domains. Mouse phenome-wide association studies revealed links to alcohol intake, nicotine response, circadian abnormalities, fat storage, glucose tolerance, and other behavioral and metabolic traits, while human analyses identified a convergent enhancer regulating *EED* and *ME3* similarly associated with substance use, abnormal sleep patterns, psychiatric traits, body weight and diabetes. This shared architecture suggests that genetic variation in pathways support energy conservation and behavioral activation may predispose individuals to SUD in modern environments^43–45^.

Our findings align with the “thrifty” hypothesis, which posits that adaptations for survival under scarcity, such as enhanced foraging and efficient fat storage, can become maladaptive in resource-rich settings^46^ and expected to operate via epigenetic mechanisms^47^. Similarly, “redox thriftiness” proposes that low mitochondrial density with high redox signaling conserves energy but promotes insulin resistance and inflammation. We find that in NPCs, the allele linked to elevated cocaine self-administration (NOD/ShiLtJ) is associated with more diffuse mitochondrial networks consistent with a metabolically thrifty state that may alter oxidative stress and synaptic plasticity^48^. Further, deletion of the enhancer region *Rr607* reduced both *Me3* expression and mitochondrial TFAM in the NOD/ShiLtJ genetic background (but not in the C57BL/6J background), suggesting a gene regulatory mechanism by which neuronal energy balance and redox signaling exert influence on brain and behavior^49^. Thus, a variant that results in a metabolically depleted state may prime mitochondria for rapid fission in striatal circuitry, increasing food seeking in times of caloric deficit but also increasing susceptibility to drug-related synaptic plasticity. Future work should dissect these mechanisms in human cell models and *in vivo*, testing how enhancer perturbation alters mitochondrial function and behavioral responses, including cocaine response, under conditions of scarcity or abundance.

Systems genetics in genetically diverse mouse populations provide a powerful framework for uncovering mechanisms that link genetic variation and vulnerability to addiction-related behaviors. Our discovery of an epigenetic mechanism connecting mitochondrial regulation to addiction-related behaviors underscores the central role of energy metabolism in neuronal plasticity. Beyond SUDs, these findings have implications for conditions such as anxiety, obesity, and sleep disturbances, which share overlapping behavioral and cellular phenotypes. The locus here identified represents one of many regulators of complex behavioral phenotypes. Elucidating these interrelated mechanisms will advance our understanding of how genetic and environmental factors converge to shape complex behaviors. Understanding these pathways will inform strategies to mitigate risk in humans and identify targets for behavioral and pharmacological intervention.

## Acknowledgements

The authors gratefully acknowledge funding from the National Institutes of Health, NIH National Institute on Drug Abuse, including R01 DA037927 to EJC, P50 DA039841 to EJC, LMT, JAB, and JDJ, U01 DA043809 to JAB, K99/R00 DA043573 to PED. We gratefully acknowledge the contribution of the JAX Data Science/Computational Sciences, Genome Technologies and Protein Sciences Services at The Jackson Laboratory for expert assistance with the work described in this publication with support from by NCI CCSG P30 CA034196. We are grateful for Miguel Skirzewski for his critical evaluation during manuscript preparation. Drug reagents were provided by the NIDA Drug Supply Program.

## Supplementary Extended Methods

### Supplementary Results

#### Behavioral trait correlations

There were 495 significant correlations among 795 unique pairwise comparisons of the behaviors tested (*padj* < 0.05, Table S3). We observed the strongest correlations, based on adjusted p-value, within behavioral assays. For example, locomotor activity in the open field assay (e.g. total distance traveled) was highly intercorrelated and measures of locomotion (e.g. percent ambulatory time and transitions) in the light-dark assay were intercorrelated (ρ = 0.28, *df* = 2730, *p* = 2.65 x 10^-60^), reflecting overall behavioral activity. We observed significant correlations across the open field and light-dark assays, for example for behavioral measures often interpreted as approach or avoidance behaviors reflective of risk-taking or anxiety-like behavior, percent time spent in the center of the open field and number of transitions in the light-dark assay, were significantly correlated (ρ = 0.21, *df* = 2730, *padj* = 1.3 x 10^-34^).

Within the IVSA procedure (Table S1), the number of sessions to acquire was negatively correlated with the number of active lever presses (ρ = -0.78, *df* = 621, *p* = 2.08 x 10^-9^) and number of infusions (ρ = -0.78, *df* = 621, *p* = 7.49 x 10^-204^) but was not correlated with any measures from the extinction and cued reinstatement phases of the protocol (Table S3).

Several cued reinstatement measures were positively correlated with extinction variables (Table S3; Figure 2A).

In the behavioral sensitization procedures for both the cocaine-treated (COC) and saline-treated (SAL) mice, we found that behavioral sensitization, indicated by the change in locomotor activity (AUC) with repeated exposure and testing from day 3 to day 11, in the COC group was significantly correlated with initial locomotor sensitivity (ρ = 0.79, *df* = 369, *padj* = 2.2 x 10^-6^), initial sensitization (ρ = 0.36, *df* = 369, *padj* = 1.7 x 10^-8^) and conditioned activation (ρ = 0.44, *df* = 369, *padj* = 6.4 x 10^-13^). However, in the SAL treated group, AUC from day 3 to day 11 was not correlated with behavioral differences observed over comparable sessions (day 2 to day 3: ρ = 0.11, *df* = 360, *padj* = 0.19; day 3 to day 5: ρ = 0.03, *df* = 360, *padj* = 0.72; day 2 to day 12: ρ = 0.05, *df* = 360, *padj* = 0.59; day 11 to day 19: ρ = 0.12, *df* = 360, *padj* = 0.16).

#### Exploration of GRM5 as possible candidate gene

The interval containing *EED* and *ME3* in the genome also includes *GRM5*, a gene previously associated with risk-taking behavior in humans (Xu, J. *et al.* 2021), and therefore its role was considered within our analyses. The enhancers regulating *EED* and *ME3* in humans are in not in linkage disequilibrium with *GRM5* (Machiela & Chanock, 2015) suggesting that fine-scale mapping is capable of discriminating between these putative loci. Moreover, in mice, eQTL data for *Grm5*^56^ does not show the allelic segregation pattern is dissociated from the traits mapped in this study. Therefore, while we note that other studies have implicated *GRM5* in mouse and human behavioral patterns consistent with those mapped in our analysis, our evidence suggests the variant acts independently of *GRM5*.

### Supplementary Tables (separate file)

Table S1. Phenotypic measures, descriptions and ontology annotations.

Table S2. Primers used for CRISPR/Cas9 mediated enhancer deletion.

Table S3. Correlation of index scores to individual measures.

Table S4. Mouse Phenome-Wide Association Study (PheWAS).

Table S5. Human PheWAS of GH11J086734 performed using the GWAS Atlas.

### Supplementary Figures

**Figure S1.**
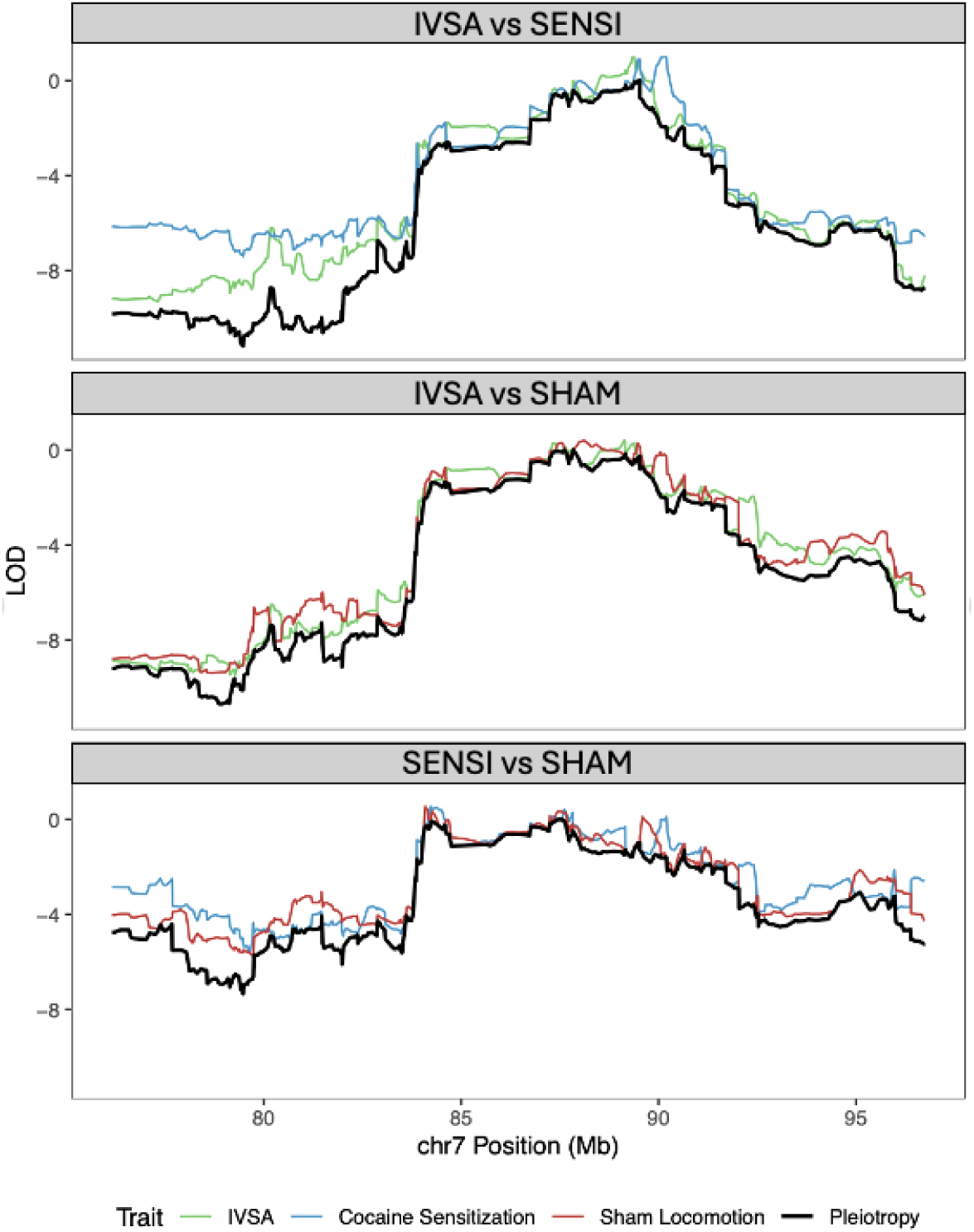
Pairwise pleiotropy analysis implicates the same locus for IVSA (indicated as green lines), COC (SENSI, blue lines), and SAL (SHAM, red lines) reference traits.

**Figure S2.**
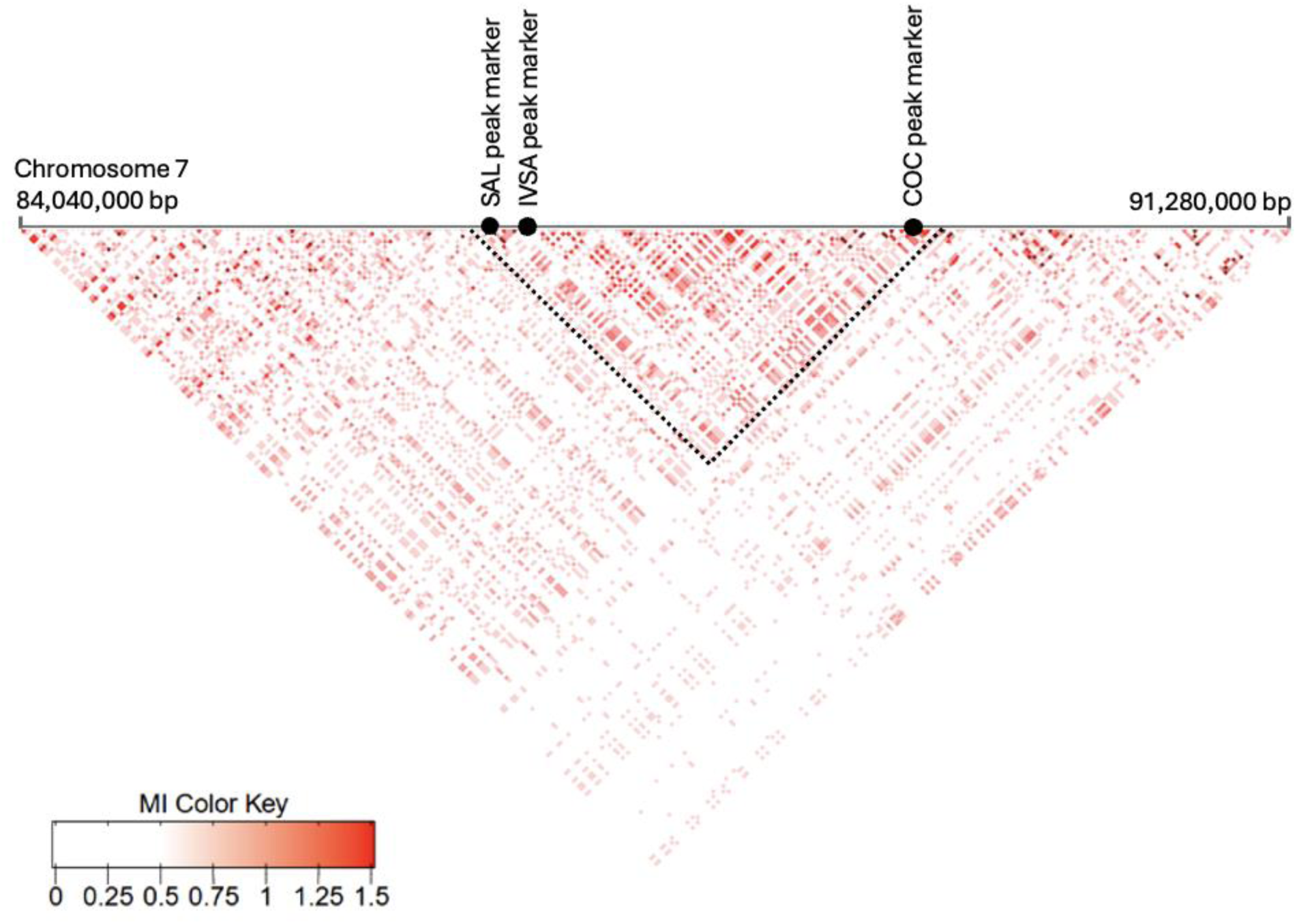
Linkage disequilibrium on chromosome 7 between 84.04 Mb to 91.28 Mb. Heatmap scale indicates mutual information (MI) at a range of zero to 1.5. We identified linkage block identified between SAL peak marker (rs220281522; chr7:87464127) and COC peak marker (rs33317667; chr7:89522474) with IVSA peak marker (rs31369570; chr7:87650445), as indicated by the grey lines.

**Figure S3.**
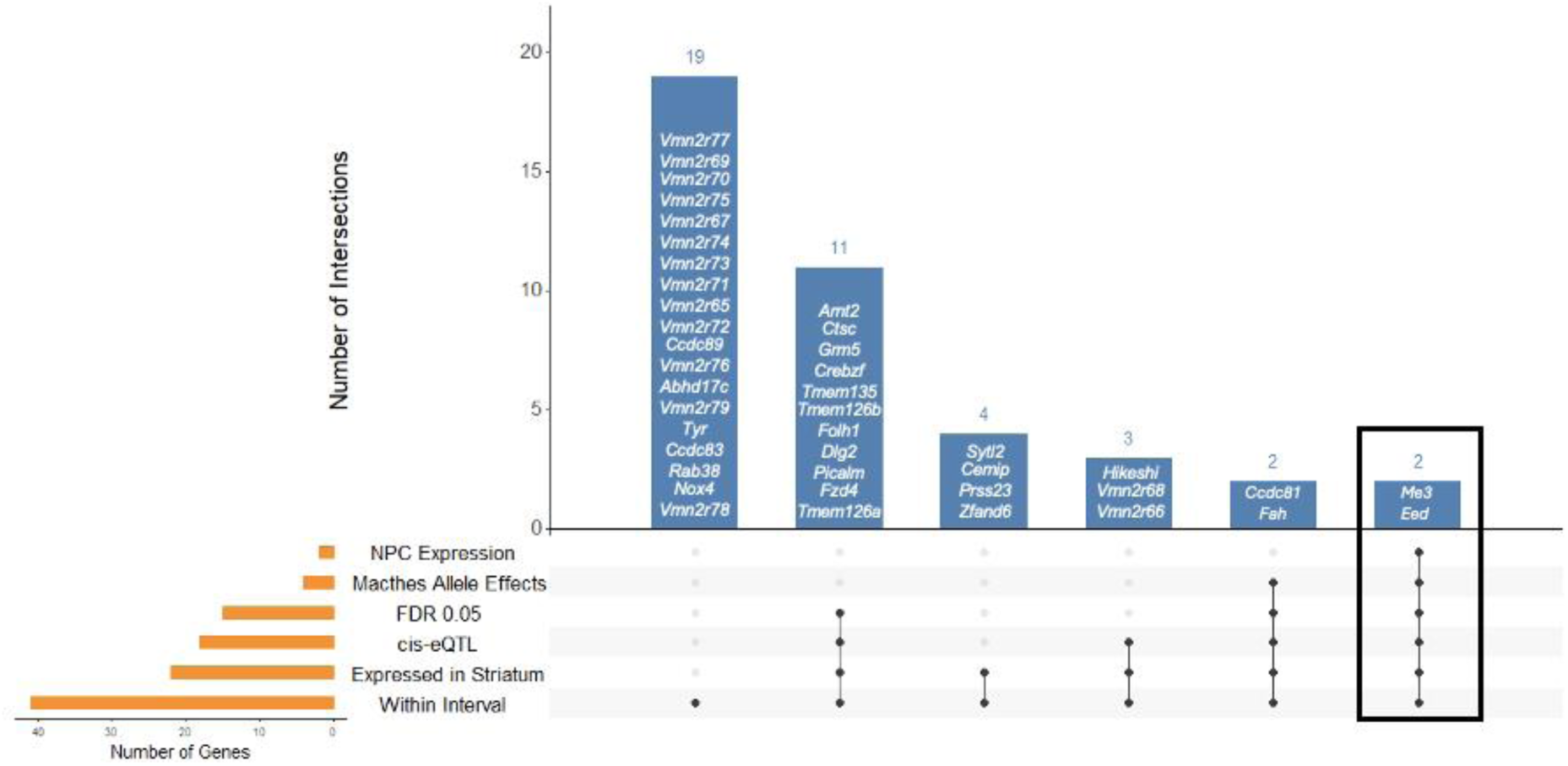
An UpSet plot of the 41 genes within the QTL interval were prioritized by filtering on those that are expressed in the striatum, and that are regulated by a significant (FDR corrected p-value < 0.05) cis eQTL with divergent C57BL/6J and NOD/ShiLtJ allele effects. The list was further refined to identify only *Me3* and *Eed* based on expression in the NPC cells, as denoted by the black box.

**Figure S4.**
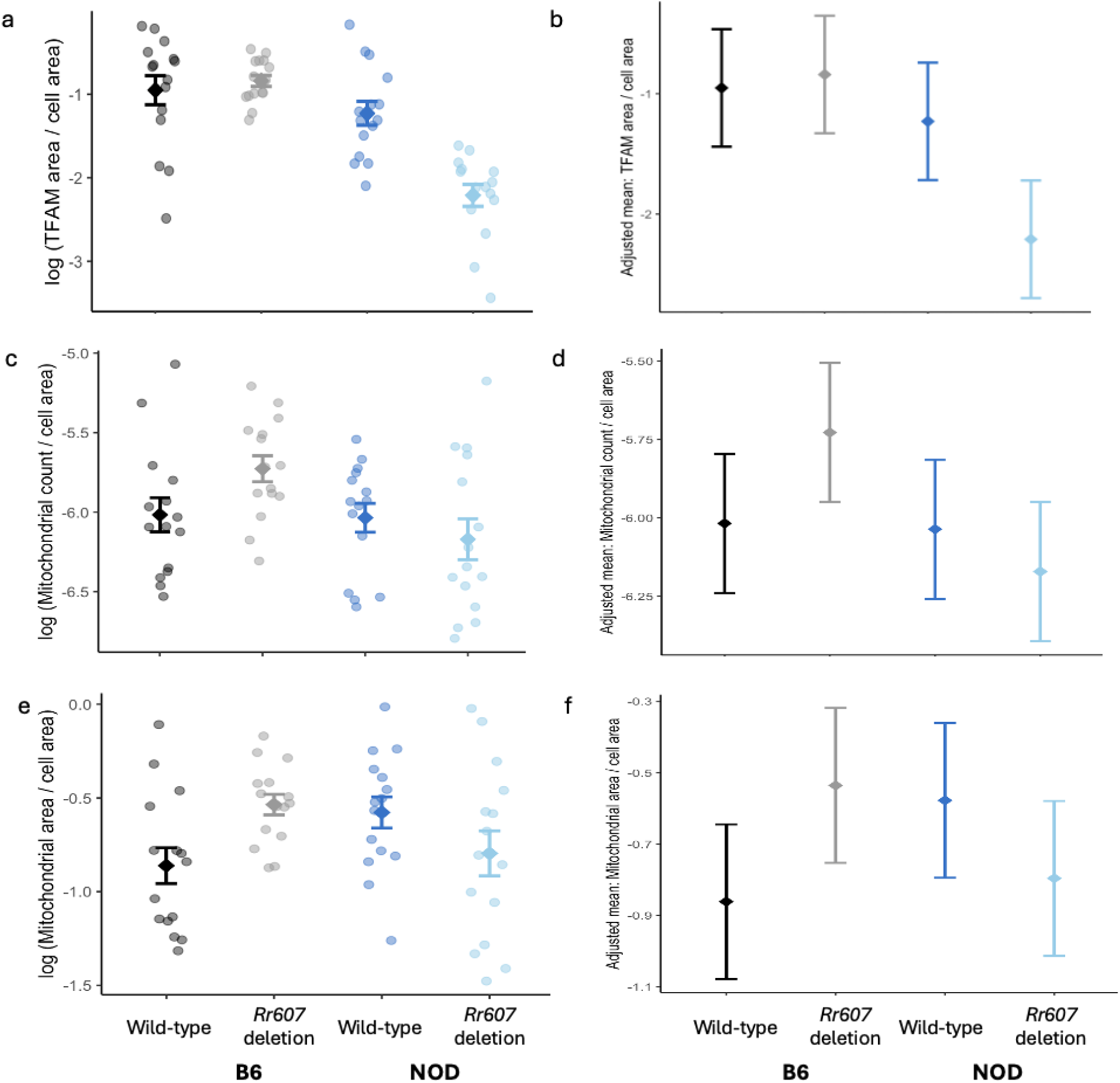
Deletion of *Rr607* enhancer induces a change in mitochondrial phenotypes in a strain-specific manner. (a) Log of TFAM area normalized to cell area and (b) corresponding linear model estimated means. (c) Log of mitochondrial count normalized to cell area and (d) model estimated means. (e) Log of mitochondrial area normalized to cell area and (f) model estimated means. Mitochondrial observed in neural progenitor cells with wild-type C57BL/6J (B6, black), C57BL/6J with *Rr607* deletion (B6, grey), wild-type NOD/ShiLtJ (NOD, dark blue), and NOD/ShiLtJ with *Rr607* deletion (NOD, light blue) backgrounds.

