## Supplemental Extended Methods for "A shared genetic regulator of metabolism and addiction-related behavior in mice and humans"

*Animals*

DO (J:DO; RRID: IMSR_MGI: 4412282) mice were obtained from The Jackson Laboratory (JAX). This genetically heterogeneous stock originates from pseudorandom mating of incipient Collaborative Cross strains, which themselves were derived from systematic intercrossing of eight inbred progenitor strains[1], collectively capture approximately 90% of the known genetic diversity within the *Mus musculus* species[2]. The DO population is maintained through continued pseudorandomized breeding at JAX[3] to generate a high density of independent recombination events, enabling high-resolution mapping of regulatory loci. For behavioral and systems genetics analyses, DO mice ($N_{males}$ = 1403, $N_{females}$ = 1496), aged 8-12 weeks and spanning generations 21 to 36, were phenotyped across the described experiments. All procedures involving vertebrate animals were approved by the Institutional Animal Care and Use Committee at JAX under protocol number 10007. Detailed information and data from all behavioral procedures can be found at the Mouse Phenome Database (accession number: CSNA03; https://phenome.jax.org/projects/CSNA03).

*Reference Trait Behavior Data Collection*

Four standard behavioral assays were conducted on consecutive days in the following order: open field, light/dark box, hole board, and novel place preference, which collectively served as “reference traits”, as all mice in the study were assessed using this battery (Figure 1).

This design enabled the projection of behavioral measures collected in subsets of mice across all experimental arms. All measured traits (Table S1) and their corresponding annotations to Vertebrate Trait[4] and Mammalian Phenotype Ontology[5] are documented in the Mouse Phenome Database[6].

Open Field (OF): To measure general exploratory activity, mice were placed in clear polycarbonate open field arenas (Med Associates- Fairfax, VT; #MED-OFAS-515U) equipped with 16 x 16 photobeam arrays at floor level and 3 inches above the floor, housed within sound-attenuating chambers (Med Associates, #MED-OFA-017). Mice were placed in the center of the arena, facing the rear wall, and tracked automatically for 60 minutes. Measurements included time spent, distance traveled, and resting duration in center, perimeter, and corners of the arena. Individuals ($N_{males}$ = 20, $N_{females}$ = 22) were excluded from OF analyses if they became sick or injured during the experiment or if there was equipment failure.

Light-Dark Box (LD): To estimate exploratory and risk-taking behavior, we used a black polycarbonate insert (Med Associates, #ENV-511) to divide the open-field arena into two equally sized compartments: a light zone and a dark zone, connected by a single access point. Mice were placed in the light zone facing the dark zone and tracked for 20 minutes. Measurements included time spent, distance traveled, and resting duration in each zone, as well as number of transitions between zones. Individuals ($N_{males}$ = 84, $N_{females}$ = 83) were excluded from LD analyses because they became sick or injured or there was an equipment failure.

Hole Board (HB): To assess exploratory and anxiety-like behavior, mice were placed in the same open field arena fitted with a hole board insert containing 16 equidistant holes (Med Associates, #ENV-515HB). Each mouse was placed in the center of the arena, facing the rear wall. Nose pokes were automatically recorded for either 10 minutes for the first cohort or 20 minutes for the second cohort. Data from the two cohorts could not be successfully combined and all hole board data were thus excluded.

Novel Place Preference (NPP): To estimate novelty seeking and response behavior, mice were tested in a three-chamber apparatus (Med Associates, #MED-CPP-3013AT) consisting of a black chamber, a white chamber, and a central gray chamber, separated by automated guillotine-style doors. Mice were initially placed in the central chamber with both doors closed for a 5-minute acclimation period. They were then randomly assigned to either the white or black chamber for a 20-minute exposure, and after which they were returned to the central chamber for an additional 5 minutes. Finally, both doors were opened, and mice were allowed to freely explore all three chambers for 10 minutes. Measurements included time spent in each chamber, resting time in the novel and central chambers, and distance traveled during this final phase. Individuals ($N_{males}$ = 105, $N_{females}$ = 108) were excluded from NPP analyses because they became sick or injured during the experiment or due to equipment failure.

*Cocaine Intravenous Self-Administration (IVSA) Data Collection*

Following reference trait testing, a subset of mice ($N_{males}$ = 537, $N_{females}$ = 546) advanced to a multi-phase cocaine intravenous self-administration testing battery as previously described[7] with minor modifications. First, mice were surgically implanted with a jugular catheter connected to a vascular access button (Instech, Plymouth Meeting, PA; #C20PU-MJV) by JAX Surgical Services (Bar Harbor, ME). After a minimum 10-day recovery period, mice then underwent a fixed-ratio FR-1 cocaine self-administration schedule training in two-lever operant chambers equipped with syringe pumps (Med Associates, #RRID:SCR_014296). After testing an initial subset of mice (cohort 1: $N_{males}$ = 44, $N_{females}$ = 36), several protocol enhancements were made to improve efficiency and maximize data collection during catheter patency for the remaining mice (cohort 2: $N_{males}$ = 493, $N_{females}$ = 510), detailed below in each IVSA phase (see *Statistical Analysis of Behavioral Phenotypes* for treatment of data from multiple cohorts).

Acquisition (AQ): During each 2-hour daily session, a press on the active lever delivered a cocaine infusion paired with a 5-second illumination of stimulus lights. This was followed by a 20-second timeout period during which no reinforcement was available. Presses of the inactive lever or the active lever during the timeout were recorded but had no programmed consequences. Mice were considered to have acquired cocaine self-administration if they received 10 or more infusions in at least 5 sessions and demonstrated less than 20% variation in the number of active lever presses across two consecutive sessions. Mice were afforded up to 28 days (cohort 1) or 18 days (cohort 2) to meet acquisition criteria, with the earliest possible completion being the fifth day of IVSA testing. 52.6% of mice reached acquisition criteria. Mice that failed to meet criteria, became sick or injured, or showed signs of lack of patency during daily catheter maintenance (e.g., solution not passing through catheter, a leaking catheter, subcutaneous pooling) were excluded from the IVSA acquisition analysis ($N_{males}$ = 235, $N_{females}$ = 244). Measurements included the number of and time between active lever presses, inactive lever presses, and infusions, in total and during the timeout period, as well as the number of sessions required to reach acquisition criteria.

Dose Response (DR): Mice that met acquisition criteria proceeded to the dose-response phase and were tested daily in 2-hour sessions with the following doses and order: 0.56, 0.32, saline, 0.18, 0.10, 0.056, 0.032, and 1.8 mg/kg/infusion (cohort 1) or 0.32, 0.10, 0.032, and 1.0 mg/kg/infusion (cohort 2). Like in the acquisition phase, an active lever press delivered a cocaine infusion and a 5-second illumination of stimulus lights, followed by a 20-second timeout period during which no reinforcement was available. Inactive lever presses and active lever presses during timeout were recorded but had no programmed consequences. Each dose was tested for up to 5 days or until stabilization was achieved, defined as less than 20% variation in the number of infusions across two consecutive sessions at the same dose. Measurements included the number of, and time between, active lever presses, inactive lever presses, and infusions, in total and during timeout period, for each dose tested.

Extinction (EX) and Reinstatement (RI): Following dose response testing, mice entered an extinction phase in which active lever presses were no longer associated with activation of the cue light, syringe pump sounds or administration of cocaine. Mice then advanced to the cued reinstatement phase, in which responses again elicited delivery of cocaine-paired cues (i.e., lights and pump sounds), but no drug was administered upon active lever press. For cohort 1, mice underwent seven daily sessions under extinction conditions, then advanced to two days of cued reinstatement testing. For cohort 2, mice underwent extinction conditions for 3 to 9 days, and advanced to cued reinstatement when the number of active lever presses fell below 50% of active lever presses on the first day of extinction and there was less than 20% variation between the final two extinction days. All cohort 2 mice advanced to cued reinstatement by day 9, regardless of whether they met extinction criteria. Mice that became sick or injured, or if there was equipment failure, were excluded from IVSA EX and RI analyses ($N_{males}$ = 43, $N_{females}$ = 46). Measurements included the number of, and time between, the formerly active lever presses and inactive lever presses.

*Cocaine Locomotor Sensitization (COC) and Saline Locomotor Behavior (SAL) Data Collection*

Following reference trait testing, a second subset of mice proceeded to a 19-day behavioral sensitization testing period to evaluate how repeated cocaine exposure alters locomotor behavior over time, as previously described[8]. Each day, mice were first habituated to the open field apparatus for 30 minutes and then injected with either saline or cocaine and returned to the open field for an additional 60 minutes, during which time locomotor activity was recorded. Mice were randomly assigned to either the cocaine sensitization group ($N_{male}$ = 191, $N_{female}$ = 185) and received saline intraperitoneal injections on days 1, 2, and 12, and cocaine injections (10 mg/kg) on days 3, 5, 7, 9, 11, and 19, or the control group ($N_{male}$ = 182, $N_{female}$ = 180) and received saline injections on all test days. Locomotor activity was quantified as total distance traveled during the 60-minute post-injection period. Behavioral variables derived from these data included: acute activation (COC) or habituation (SAL) from day 2 to day 3, initial sensitization (COC) or habituation (SAL) from day 3 to day 5, conditioned activation (COC) or habituation (SAL) from day 2 to day 12, sensitization expression (COC) or habituation (SAL) from day 11 to day 19, COC five-trial sensitization (area under the curve; AUC) or SAL activity AUC from day 3 to day 11.

*Statistical Analysis of Behavioral Phenotypes*

All data processing and statistical analyses were conducted in R (Version 4.1.2) [9]. The full analysis code is publicly available at: 10.5281/zenodo.17298531.

We calculated correlations among all reference traits, among IVSA phenotypes and reference traits, COC phenotypes and reference traits, and among SAL phenotypes and reference traits (N = 795 correlations) using the *cor.test* function and Spearman's method in R, then applied the *p.adjust* function using the Benjamini-Hochberg method[10] to control for false discovery rate.

For data collected across the two cohorts, if we observed differences in the mean and variance of measured variables, we addressed these effects using rank-normal transformations using a custom *norm_rank_transform* function in R separately within cohort 1 and cohort 2 variables, then reassessed mean and variance. In only one case did the mean and variance continue to differ between cohorts: measures of the hole board assay. Therefore, we excluded hole board data from downstream analyses to avoid introducing bias.

*Canonical Correlation*

We applied the reference trait genetics framework we previously described in Skelley et al [11] to identify biological and behavioral predictors of cocaine self-administration and behavioral sensitization using the partially disjoint datasets that resulted from our testing protocol. This approach uses canonical correlation analysis and the *cancor* function in R, to relate a linear combination of reference traits measured in all mice to “target traits” (i.e., traits measured in IVSA, COC, and SAL subsets of mice). This enables the imputation of target trait values for mice not directly tested in those behavioral paradigms. To minimize overfitting, we implemented a forward variable selection strategy into canonical correlations[12]. We began by selecting key addiction-related traits: AUC for behavioral sensitization and active lever presses during cued reinstatement for IVSA. We then identified the novelty response and risk-taking reference traits most strongly correlated with each target trait to serve as the initial variables for canonical correlation analysis. Subsequently, we evaluated the inclusion of additional variables using a permutation-based significance test. For each candidate variable, we performed 1,000 permutations of random shuffling of the novelty response variables, which conserves correlation between novelty response and target trait sets while randomizing the relationship between the reference and target trait. Randomization was accomplished across subjects using a bootstrapping loop with the *sample* function in R. After each permutation, we calculated the magnitude of the first canonical correlate to generate a null distribution. A variable was retained if its inclusion resulted in a canonical correlation exceeding the $99.99^{th}$ percentile of the null distribution ($p < 0.001$). We required a minimum of 300 observations across reference and target traits and limited the final model to a maximum of 20 variables to further guard against overfitting. The final canonical correlates were projected onto the full DO cohort using linear regression *lm* function in R (N = 2,682 mice), generating imputed scores for IVSA, COC, and SAL for each mouse.

*Genotyping*

Genomic DNA was isolated from tail samples and genotyped using the Giga Mouse Universal Genotyping Array (GigaMUGA) at NEOGEN (http://www.neogen.com/GeneSeek). This array is based on the Illumina Infinium II platform and includes approximately 143,000 single nucleotide polymorphism (SNP) markers optimized for high-resolution haplotype reconstruction in multi-founder populations such as the DO mice. Of these, 54,250 SNPs were selected to maximize informativeness in the DO population, removing SNP genotypes that do not fully define the corresponding founder haplotype. The resultant array achieves a marker density of approximately one informative SNP per 100 kilobases, enabling fine-scale mapping[13].

*QTL Mapping and Pleiotropy Analysis*

To identify regions of the genome associated with the imputed scores for each target trait—IVSA, COC, and SAL— we preformed QTL analysis using the R/qtl2 package (RRID:SCR_018181)[14]. Quality control was performed on raw genotype data within the R/qtl2 package. We used the *drop_nullmarkers* function to omit markers without any genotype data and *n_missing* function to identify samples missing data (low call rates). Additionally, we compared sex in the metadata with X and Y chromosome genotypes and removed sample duplicates. Our QTL mapping model included additive covariates for DO generation and sex. Genome-wide significance thresholds were determined using 1,000 permutations using the *scan1perm* function from R/qtl2 to estimate the 5% false discovery rate. For each significant QTL, 1.5 LOD intervals were defined around each peak and combined to define the positional candidate region of the pleiotropic regulatory locus. Additionally, we extracted the 8-state founder allele effect coefficients for downstream analyses, using *calc.genoprob* function in the R/qtl2 package. To assess pleiotropy, we used the *scan-pvl_2* function R/qtl2pleio package[15].

Specifically, we examined all loci within ±10 Mb of the QTL peaks for IVSA, COC and SAL scores to identify overlapping signals indicative of shared regulatory mechanisms. All code is available at 10.5281/zenodo.17298531

*Candidate Gene Prioritization in QTL Interval*

To prioritize candidate genes within each QTL interval, we ranked genes based on their relevance to each of the three target traits (IVSA, COC and SAL) using the following criteria: (1) cis-eQTL significance and allelic effect concordance: Genes with significant cis-eQTLs whose allelic segregation patterns matched those observed at the peak behavioral QTL were prioritized; (2) Log of odds (LOD) score correlation: we assessed the strength of correlation between the LOD scores of all variants within 10 Mb window surrounding either the eQTL peak or the behavioral QTL peak; and (3) founder coefficient correlation: we calculated the absolute correlation between the 8-state founder allele effect coefficients from the R/qtl2 mapping model at the peak eQTL marker and those at the peak behavioral QTL marker. Each gene was ranked independently on each of these three criteria. The final prioritization score was computed as the average of the three individual ranks, yielding a composite rank for each candidate gene.

*Linkage disequilibrium*

After quality control on raw genotype data by R/qtl2 package, genotype data were formatted into Plink (RRID:SCR_001757)[16] file format as inputs for efficient linkage disequilibrium (LD) calculation. Compared with traditional LD measures (D/D′ and $r2$), the mutual information (MI), has been used for multilocus haplotype LD and epistasis analyses, was calculated in a parallelized way by R/GWLD package[17]. The calculated LD results of MI values for chromosome 7 region from 84.04 Mb to 91.28 Mb were then visualized as LD heatmap to show LD status on peak marks. All code is available at 10.5281/zenodo.1729853.1

*Striatal Single-Nucleus Multiomic Profiling in DO Founder Strains*

To understand how genetic variation within the DO population affects gene regulation and cellular function in the striatum, we estimated gene expression and mapped chromatin accessibility using single-nucleus RNA sequencing (snRNA-Seq) and ATAC-seq simultaneously in the same cell, which enables detection of cell-type specific cis-regulatory elements, resulting in increased sensitivity and specificity of enhancer identification[18]. We used our publicly available dataset through the Gene Expression Omnibus (GEO, RRID:SCR_005012) under accession number GSE228530[18]. The dataset included striatal tissue that had been collected from two male and two female 12-week old, drug-naïve mice that had undergone behavioral phenotyping, of each of the eight DO founder strains: A/J (RRID: MGI:2159747), 129S1/SvImJ (RRID: MGI:2671655), CAST/EiJ (RRID: MGI:2160389), C57BL/6J (RRID: MGI:3028467), NOD/ShiLtJ (RRID: MGI:2159775), NZO/HlLtJ (RRID: MGI:2668669), PWK/PhJ (RRID: MGI:2159876), and WSB/EiJ (RRID: MGI:2160618).

Downstream gene expression analysis was performed using Seurat (version 4.0.3; RRID:SCR_007322)[19]. Low-quality cells (nuclei) and multiplets were filtered out by excluding cells with fewer than 200 or more than 7,500 detected genes, or with >15% mitochondrial gene content[18]. Cell clusters were annotated using marker genes identified through the *FindAllMarkers* function and compared to reference annotations from the DropViz mouse striatum dataset[20]. Between-strain expression differences were assessed using two-tailed two-sample t-tests using the *distuv.studentsT* function in the *distuv* package, then applied the *p.adjust* function using the Benjamini-Hochberg method to control for false discovery rate. Enhancer-gene links were identified using the Enhlink package[18]. Significance of inferred links were determined using a one-sample t-test on 100 bootstrap iterations (df = 99) to test if the mean information gain score was greater than zero. P-values were then corrected for multiple comparisons using the Benjamini-Hochberg method.

*Validation of Rr607 as a regulator of Me3/Eed expression*

To confirm the effect of the putative enhancer, *Rr607* (Peak A*;* chr7:89837035–89837873 [839bp]) on gene expression regulation and expected cellular characteristics, we used previously established mouse embryonic stem cell (mESC) lines from C57BL/6J (B6 4-1) and NOD/ShiLtJ (AC576) backgrounds[21,22] (RRID:CVCL_2H72) for CRISPR/Cas9-mediated genome editing. Cells were cultured in standard mESC media consisting of Dulbecco's Modified Eagle's Medium (DMEM; GIBCO #11960069), 15% Fetal Bovine Serum (FSB; GIBCO #10439024), 2 mM GlutaMAX (GIBCO #35050061), 50 U/ml Penicillin-Streptomycin (GIBCO #15140122), 1X Non-Essential Amino Acids (GIBCO #11140050), 1 mM Sodium Pyruvate (GIBCO #11360070), 55 µM 2-Mercaptoethanol (GIBCO #21985023), 1,000 IU of Leukemia Inhibitory Factor (LIF, R&D Systems #8878-LF-500), 1 µM PD0325901 (StemCell Technologies #72184), and 3 µM CHIR99021 (StemCell Techonologies #72054). Unless otherwise specified, cells were grown on gelatinized (StemCell Technologies #07903), tissue culture-coated plasticware at 37°C in a humidified incubator with 5% $CO_2$. Media were replenished daily.

To determine the effect of the *Rr607* (Peak A) region on *Me3* and *Eed* gene expression, genome editing was used to generate a homozygous deletion of a putative enhancer region regulating *Me3 and Eed* in mESC lines derived from C57BL/6J and NOD/ShiLtJ backgrounds, representing low and high allele effects and were easily amendable to genomic engineering. All guide RNA and primer sequences were designed using the C57BL/6J reference genome (GRCm38/mm10). Candidate target regions were selected based on the absence of polymorphisms in both C57BL/6J and NOD/ShiLtJ strains, as confirmed using publicly available variant databases such as GenomeMUSter[23] (RRID:SCR_024214).

Guide RNAs were designed to flank the targeted enhancer region, *Rr607,* at chr7:89837084–89837103 and chr7:89837833–89837852 (GRCm38/mm10; Table S2**).** Sequence-specific CRISPR RNAs (crRNAs) were synthesized by IDT and resuspended to 100µM in nuclease-free duplex buffer (IDT #11-01-03-01). To form guide duplexes, 1 µl of crRNA was combined with 1 µl of 100uM TracrRNA (IDT #1075927) in 98 µl of nuclease-free duplex buffer and heated at 95°C for 5 minutes. For transfection in a 12-well plate, the ribonucleoprotein (RNP) complex was assembled by combining 171 µl of Opti-MEM (GIBCO #51985034) with 6 µl each of upstream and downstream guide duplexes, 12 µl of 1 µM Cas9 enzyme (IDT #1081058), and 5 µl of Cas9+ reagent (Invitrogen #CMAX00001). This mixture was added to 192 µl of Opti-MEM and 8 µl of CRISPRMAX transfection reagent (Invitrogen #CMAX00001), then incubated at room temperature for 20 minutes. During incubation, mESCs were dissociated, counted, and diluted to 400,000 cells/ml in antibiotic-free culture medium. A total of 1 ml of cells was combined with 400 µl of transfection complex and plated into one well of a 12-well plate. After 24 hours, cells were sorted for ATO550 fluorescence using a FACSymphony S6 (RRID:SCR_022538), and ATO550-positive cells were plated at 250 cells/ml onto 100mm culture dishes pre-seeded with mouse embryonic fibroblasts in standard mESC medium.

After 5-7 days, individual colonies were picked, expanded, and screened for the deletion event by PCR using primers listed in Table S2. A total of 96 clones per editing event were screened by agarose gel electrophoresis, and those showing evidence of deletion were confirmed by Sanger sequencing[24]. Two to three homozygous deletion clones were expanded per editing event, and one validated clone per strain was selected for downstream experiments. The final edited mESC lines used in follow-up studies were: C57BL/6J deletion—C57BL/6J Peak A (C57BL/6J-Rr607$^{em1Ejc}$/Ejc), clone B4; C57BL/6J control—B6 scramble (wild-type), clone

A4; NOD/ShiLtJ deletion—NOD Peak A (NOD/ShiLtJ-Rr607$^{em2Ejc}$/Ejc), clone F6; and NOD/ShiLtJ control—NOD scramble (wild-type), clone B7.

Mouse embryonic stem cells were differentiated into NPCs as previously described[25]. For each genome-edited mESC line, differentiation was performed in triplicate to generate three independently derived NPC lines. NPC were maintained in NPC Maintenance Media (NMM)[25], consisting of 1:1 v/v Neurobasal (GIBCO #21103049) and DMEM/F12 (GIBCO #10565018), 0.5X N2 supplement (GIBCO #17502048), 0.5X B27 supplement (GIBCO #17504001), 5 µg/ml insulin (Millipore #4512-01), 1mM L-Glutamine (GIBCO #A2916801), 50 U/ml Penicillin-Streptomycin (GIBCO #15140122), 0.5x NEAA (GIBCO #11140050), and 27.5 µM (GIBCO #21985023). Cells were cultured on TC-coated plasticware treated with 5 µg/ml fibronectin (Fn, Millipore #FC010), 1 µg/ml laminin (Sigma-Aldrich #L2020), and 100 µg/ml poly-L-lysine (Sigma-Aldrich #P8920). Cultures were maintained at 37°C in a humidified incubator with 5% $CO_2$ and fed daily with NMM.

To assess the impact of enhancer deletion on the expression of *Eed*, *Me3*, and *Ccdc81* and markers of neuronal differentiation, RNAseq was performed on NPCs derived from three biological replicates of each edited and control mESC line. NPCs were thawed onto PLL/Fn/Lm-coated dishes and cultured in NMM for one passage. Upon reaching confluency, cells were dissociated using TrypLE (GIBCO #12605010), rinsed with PBS (GIBCO #20012050), pelleted, and snap frozen on dry ice. Total RNA was extracted from $1x10^6$ cells using the NucleoMag RNA Kit (Macherey-Nagel) on the KingFisher Flex purification system (ThermoFisher), following the manufacturer's instructions. Cells were lysed in MR1 buffer (Macherey-Nagel) by vortexing.

RNA concentration and quality were assessed using the Nanodrop 2000 spectrophotometer (Thermo Scientific; RRID:SCR_018042) and the RNA ScreenTape Assay (Agilent Technologies). All samples had 260/280 ratios between 1.95 – 2.01, RNA Integrity Numbers (RINs) > 8.0, and 28s/18s ratios > 1.0 (range: 9.7 – 10 RIN). Stranded mRNA libraries were constructed using the KAPA mRNA HyperPrep Kit (Roche Sequencing and Life Science), following the manufacturer's instructions. Briefly, poly(A)+ mRNA was isolated using oligo-dT magnetic beads, followed by RNA fragmentation, first- and second-strand cDNA synthesis, ligation of Illumina-specific adapters with unique barcodes, sequence for each library, and PCR amplification. Library quality and concentration were assessed using the D5000 ScreenTape assay (Agilent Technologies) and Qubit dsDNA HS Assay (ThermoFisher), respectively. Libraries were sequenced 150 bp paired-end reads on an Illumina NovaSeq X Plus platform (RRID:SCR_024568) using the 10B Reagent Kit. Each sample yielded >300,000 reads (range 333,000 – 338,000 reads).

To estimate differential gene expression from the NPC samples, reads were generated from raw data and demultiplexed using BCL2Fastq (v2.18.0.12) concatenated by sample and aligned using the GBRS method[26] which accounts for strain-specific read alignment and incorporates the GRCm38 reference assembly. Differential gene expression analysis was performed using the DESeq2 package (version 1.48.1; Love *et al.*, 2014) in R (version 4.5). Raw read counts obtained from RNA-seq were first imported and organized into a DESeqDataSet object, with experimental design factors specified according to treatment groups. Genes with low counts across all samples were filtered out prior to normalization. DESeq2's median-of-ratios method was applied to normalize for library size and compositional bias. Differential expression was estimated using a negative binomial generalized linear model, with Wald tests[27] or likelihood ratio tests[28] (as appropriate) employed to assess statistical significance. Resulting $p$-values were adjusted for multiple testing using the Benjamini–Hochberg false discovery rate correction. Genes with an adjusted $p < 0.05$ and $|\log_2$ fold change$| \geq 1$ were considered significantly differentially expressed. Normalized expression values were transformed using the variance stabilizing transformation or visualization and downstream analyses, including principal component analysis and clustering.

*Immunofluorescence imaging of NPC mitochondria*

To determine whether the regulatory locus has downstream effects on mitochondrial structure or abundance, NPCs derived from three biological replicates of each enhancer-deletion and control clone were thawed onto PLL/Fn/Lm-coated culture dishes and maintained in 24-well plate NMM for one passage. Upon reaching ~80% confluency, cells were dissociated with TrpLE, washed in PBS pelleted, and plated at 500,000 cells per well in NMM onto 14 mm glass PLL/Fn/Lm-coated coverslips, fixed with 4% paraformaldehyde for 20 minutes at room temperature. Cells were then permeabilized with 0.1% Triton X-100 (ThermoFisher Scientific, #BP151-100) for 5 minutes and blocked with PBS containing 5% fetal bovine serum (VWR, #97068-085) for 30 minutes. Immunostaining was performed using the following primary antibodies: anti-TFAM (RRID:AB 1182588, Proteintech, #22586-1-AP) and anti-HSP60 (RRID:AB_631683, Santa Cruz Biotechnology, #sc-1052), incubated for 1 hour at room temperature. After washing, cells were incubated with secondary antibodies for 1 hour: rhodamine red donkey anti-rabbit (RRID:AB_2340613, Jackson ImmunoResearch, #711-295-152) and Alexa Flour 647 bovine anti-goat (RRID:AB_2340885, Jackson ImmunoResearch, #805-605-180). Nuclei were counterstained with 200 ng/ml DAPI (ThermoFisher Scientific, #62247) for 2 minutes. Washes were performed between each step, and coverslips were mounted using ProLong Diamond Antifade Mountant (Invitrogen, #MP33025). Z-stack images (five fields of view from each of three biological replicates per condition) were acquired using a 60X oil immersion objective on a Nikon ECLIPSE Ti2 widefield microscope. Maximum intensity projections of each image were generated using NIS-Elements AR 5.21.02 (RRID:SCR_027181) software. Mitochondrial network analysis was then completed in ImageJ (RRID:SCR_003070) by creating a binary mask on the HSP60 images, then the Mitochondria Analyzer tool (RRID:SCR_027707)[29] was used to calculate the mitochondrial count and total mitochondrial area for each image per biological replicate. For statistical analysis, these measurements were normalized to the total cellular area in each field of view as determined by drawing regions of interest around each cell. We then created a binary mask over TFAM, converted the mask to a region of interest, and utilized the measurement tool in ImageJ to determine TFAM area. Similar to mitochondrial count and area, we normalized TFAM area to cell area for analyses. In R[9] (version 4.5.2), we log transformed all normalized variables and fit a separate linear mixed-effect model for each using the *lmer* function in the lme4 package to test the fixed effects of strain (C57BL/6J or NOD/ShiLtJ), genotype (wild-type or *Rr607* deletion) and their interaction, with the random effect of biological replicate nested within trait. Pairwise contrasts between strain and genotype combinations were computed using the *emmeans* function in the emmens package, and plotted using the ggplot2 package in R.

*Bioinformatics*

To identify additional mouse phenotypes beyond those measured in this study and also associated with *Rr607* mouse enhancer, we first identified variants that are in linkage disequilibrium (LD) with the six variants used to map the concensus QTL to chr7: 89837366 – 89837866 (GRCm38) : rs31660604, rs32308187, rs31326011, rs31051447, rs31887897, rs49826726. GenomeMUSter[23] (RRID:SCR_024214) allelic state data for 657 mouse strains provided the dense genetic backdrop necessary to estimate LD. D-prime (d’) was calculated pairwise between each of the six variants and the set of variants in the Mouse Phenome Database [30] (RRID:SCR_003212 MPD) Trait Regulatory Network within 500KB of the region, chr7: 89337366 – 90337866 (Supplemental Code: calculate_ld_diploid.R). Variants in LD (d’ ≥ 0.80) with at least one of the six variants were included in downstream analysis. If the allelic state in GenomeMUSter was different between NOD/ShiLtJ and C57BL/6J, the variant was annotated as ‘different_allele = 1’. Traits regulated by variants in LD with at least one of the six variants of interest were included if the GWAS variant effect had a p-value ≤ 0.01. Strain means of traits were downloaded from MPD (phenome.jax.org/downloads - accessed on Oct 30 2025). If measured, the mean z-score for NOD/ShiLtJ and C57BL/6J are provided (NOD/ShiLtJ_z, C57BL/6J_z) as well as the mean difference of NOD/ShiLtJ in comparison to C57BL/6J. Trait measurement details and ontology information were merged with the regulated traits, downloaded from MPD (phenome.jax.org/downloads – accessed on Oct 30, 2025). For each trait and each of the six variants, we summarized the results by calculating the mean d' and mean -log10p across variants that regulate the trait and were in LD with the variant. The number of variants ('n') and the number of variants with different allelic states ('num_variants_w_different_alleles') were included for each trait.

To investigate whether a gene regulatory mechanism involving *Eed* and *Me3* is associated with SUD-related phenotypes across species, we queried these genes in GeneWeaver's multi-species database (RRID:SCR_009202, 3/27/2025), which integrates over 300,000 gene sets derived from curated annotations, published studies, archived experiments, and other sources[31]. The results of this query were filtered to identify gene sets relevant to SUDs, specifically those associated with acute or chronic exposure to, or self-administration of, drugs of any kind (eg., cocaine, opioids, alcohol, nicotine, etc.).

To evaluate whether a regulatory mechanism exists in humans similar to *Rr607* (i.e., Peak A) in mice that jointly regulates *Eed* and *Me3,* we queried the GeneHancer[32] database (RRID:SCR_023953, 6/19/2025) within GeneCards Version 5.24 (RRID:SCR_002773). We identified three enhancers, GH11J086589 located at chr11:86589776-86597975 (GRCh38/hg38), GH11J086475 located at chr11:86475897-86478555, and GH11J086734 located at chr11:86734911-86749108 that each influence the expression of *EED* and *ME3*; GH11J086589 and GH11J086475 also regulates the expression of *CCDC81*. We next prioritized variants within the chromosomal regions associated with each of the enhancers by quering the dbSNP database[33] (RRID:SCR_002338, 6/19/2025) within RegulomeDB[34] (RRID:SCR_017905, 6/19/2025), and identifed 145 variants (GH11J086589 N = 56; GH11J086475 N = 11; GH11J086734 N = 75), 13 of which we prioritized their Regulome rank of '1a' or '1b' based amount of supporting evidence from eQTL, chromatin accessiblity peak, transcription factor binding sites, motif and footprint sources. We further focused on

GH11J086734, and specifically the 7 prioritized variants (Regulome 1a = rs17149343; Regulome 1b = rs17758206, rs2433439, rs10898531, rs2433436, rs2433432, rs71465654) because we found that deletion of the enhancer *Rr607* (Peak A) in mouse neural progentor cells did not alter expression of *Ccdc81,* therefore the enhacer in humans that is not associated with *CCDC81,* GH11J086734, is the most similiar to the mouse *Rr607* enhancer.

To identify human phenotypes associated with the GH11J086734 enhancer, which functions similiarly to the *Rr607* mouse enhancer, we queried GWAS Atlas[35] for the 7 prioritized variants within GH11J086734, as well as rs10501688, an intronic variant in the nearby gene, GRM5, that has been associated with risk-taking behavior [36] GWAS Atlas preforms a Phenome-Wide Association Study (PheWAS) analysis thresholded at p-value < 0.05, and reports phenotypes associated for each rsID which we used to evaluate the variants' association with psychiatric, metobolic, and neurological phenotypes.


1 Svenson, K. L. *et al*. High-resolution genetic mapping using the Mouse Diversity outbred population. *Genetics* **190**, 437–447 (2012). https://doi.org/10.1534/genetics.111.132597

2 Roberts, A., Pardo-Manuel de Villena, F., Wang, W., McMillan, L. & Threadgill, D. W. The polymorphism architecture of mouse genetic resources elucidated using genome-wide resequencing data: implications for QTL discovery and systems genetics. *Mamm Genome* **18**, 473–481 (2007). https://doi.org/10.1007/s00335-007-9045-1

3 Chesler, E. J. *et al*. Diversity Outbred Mice at 21: Maintaining Allelic Variation in the Face of Selection. *G3 (Bethesda)* **6**, 3893–3902 (2016). https://doi.org/10.1534/g3.116.035527

4 Park, C. A. *et al*. The Vertebrate Trait Ontology: a controlled vocabulary for the annotation of trait data across species. *J Biomed Semantics* **4**, 13 (2013). https://doi.org/10.1186/2041-1480-4-13

5 Smith, C. L. & Eppig, J. T. Expanding the mammalian phenotype ontology to support automated exchange of high throughput mouse phenotyping data generated by large-scale mouse knockout screens. *J Biomed Semantics* **6**, 11 (2015). https://doi.org/10.1186/s13326-015-0009-1

6 (!!! INVALID CITATION !!! 35).

7 Dickson, P. E. *et al*. Systems genetics of intravenous cocaine self-administration in the BXD recombinant inbred mouse panel. *Psychopharmacology (Berl)* **233**, 701–714 (2016). https://doi.org/10.1007/s00213-015-4147-z
8 Schoenrock, S. A. *et al*. The collaborative cross strains and their founders vary widely in cocaine-induced behavioral sensitization. *Front Behav Neurosci* **16**, 886524 (2022). https://doi.org/10.3389/fnbeh.2022.886524
9 R: A Language and Environment for Statistical Computing v. 4.1.2 (R Foundation for Statistical Computing, Vienna, Austria, 2025).
10 Benjamini, Y. & Hochberg, Y. Controlling the False Discovery Rate: A Practical and Powerful Approach to Multiple Testing. *Journal of the Royal Statistical Society: Series B (Methodological)* **57**, 289–300 (2018). https://doi.org/10.1111/j.2517-6161.1995.tb02031.x
11 Skelly, D. A., Raghupathy, N., Robledo, R. F., Graber, J. H. & Chesler, E. J. Reference Trait Analysis Reveals Correlations Between Gene Expression and Quantitative Traits in Disjoint Samples. *Genetics* **212**, 919–929 (2019). https://doi.org/10.1534/genetics.118.301865
12 Winkler, A. M., Renaud, O., Smith, S. M. & Nichols, T. E. Permutation inference for canonical correlation analysis. *Neuroimage* **220**, 117065 (2020). https://doi.org/10.1016/j.neuroimage.2020.117065
13 Morgan, A. P. *et al*. The Mouse Universal Genotyping Array: From Substrains to Subspecies. *G3 (Bethesda)* **6**, 263–279 (2015). https://doi.org/10.1534/g3.115.022087
14 Broman, K. W. *et al*. R/qtl2: Software for Mapping Quantitative Trait Loci with High-Dimensional Data and Multiparent Populations. *Genetics* **211**, 495–502 (2019). https://doi.org/10.1534/genetics.118.301595
15 Boehm, F. J., Chesler, E. J., Yandell, B. S. & Broman, K. W. Testing Pleiotropy vs. Separate QTL in Multiparental Populations. *G3 (Bethesda)* **9**, 2317–2324 (2019). https://doi.org/10.1534/g3.119.400098
16 Purcell, S. *et al*. PLINK: a tool set for whole-genome association and population-based linkage analyses. *Am J Hum Genet* **81**, 559–575 (2007). https://doi.org/10.1086/519795
17 Zhang, R. *et al*. GWLD: an R package for genome-wide linkage disequilibrium analysis. *G3 (Bethesda)* **13** (2023). https://doi.org/10.1093/g3journal/jkad154
18 Poirion, O. B. *et al*. Enhlink infers distal and context-specific enhancer-promoter linkages. *Genome Biol* **25**, 235 (2024). https://doi.org/10.1186/s13059-024-03374-9
19 Hao, Y. *et al*. Integrated analysis of multimodal single-cell data. *Cell* **184**, 3573–3587 e3529 (2021). https://doi.org/10.1016/j.cell.2021.04.048
20 Saunders, A. *et al*. Molecular Diversity and Specializations among the Cells of the Adult Mouse Brain. *Cell* **174**, 1015–1030 e1016 (2018). https://doi.org/10.1016/j.cell.2018.07.028
21 Czechanski, A. *et al*. Derivation and characterization of mouse embryonic stem cells from permissive and nonpermissive strains. *Nat Protoc* **9**, 559–574 (2014). https://doi.org/10.1038/nprot.2014.030

22 Skelly, D. A. *et al*. Mapping the Effects of Genetic Variation on Chromatin State and Gene Expression Reveals Loci That Control Ground State Pluripotency. *Cell Stem Cell* **27**, 459–469 e458 (2020). https://doi.org/10.1016/j.stem.2020.07.005
23 Ball, R. L. *et al*. GenomeMUSter mouse genetic variation service enables multitrait, multipopulation data integration and analysis. *Genome Res* **34**, 145–159 (2024). https://doi.org/10.1101/gr.278157.123
24 Sanger, F., Nicklen, S. & Coulson, A. R. DNA sequencing with chain-terminating inhibitors. *Proc Natl Acad Sci U S A* **74**, 5463–5467 (1977). https://doi.org/10.1073/pnas.74.12.5463
25 Cortes, D. E. *et al*. An in vitro neurogenetics platform for precision disease modeling in the mouse. *Sci Adv* **10**, eadj9305 (2024). https://doi.org/10.1126/sciadv.adj9305
26 Choi, K. *et al*. Genotype-free individual genome reconstruction of Multiparental Population Models by RNA sequencing data. *bioRxiv*, 2020.2010.2011.335323 (2025). https://doi.org/10.1101/2020.10.11.335323
27 Wald, A. Tests of statistical hypotheses concerning several parameters when the number of observations is large. *Trans. Amer. Math. Soc* **54**, 426–482 (1943).
28 Neyman, J. & Peason, E. On the Use and Interpretation of Certain Test Criteria for Purposes of Statistical Inference. *Biometrika* **20A**, 175–240 (1928).
29 Chaudhry, A., Shi, R. & Luciani, D. S. A pipeline for multidimensional confocal analysis of mitochondrial morphology, function, and dynamics in pancreatic beta-cells. *Am J Physiol Endocrinol Metab* **318**, E87–E101 (2020). https://doi.org/10.1152/ajpendo.00457.2019
30 Bogue, M. A. *et al*. Mouse phenome database: curated data repository with interactive multi-population and multi-trait analyses. *Mamm Genome* **34**, 509–519 (2023). https://doi.org/10.1007/s00335-023-10014-3
31 Baker, E., Bubier, J. A., Reynolds, T., Langston, M. A. & Chesler, E. J. GeneWeaver: data driven alignment of cross-species genomics in biology and disease. *Nucleic Acids Res* **44**, D555–559 (2016). https://doi.org/10.1093/nar/gkv1329
32 Fishilevich, S. *et al*. GeneHancer: genome-wide integration of enhancers and target genes in GeneCards. *Database (Oxford)* **2017** (2017). https://doi.org/10.1093/database/bax028
33 Sherry, S. T. *et al*. dbSNP: the NCBI database of genetic variation. *Nucleic Acids Res* **29**, 308–311 (2001). https://doi.org/10.1093/nar/29.1.308
34 Boyle, A. P. *et al*. Annotation of functional variation in personal genomes using RegulomeDB. *Genome Res* **22**, 1790–1797 (2012). https://doi.org/10.1101/gr.137323.112
35 Watanabe, K. *et al*. Author Correction: A global overview of pleiotropy and genetic architecture in complex traits. *Nat Genet* **52**, 353 (2020). https://doi.org/10.1038/s41588-019-0571-z
36 Baselmans, B. *et al*. The Genetic and Neural Substrates of Externalizing Behavior. *Biol Psychiatry Glob Open Sci* **2**, 389–399 (2022). https://doi.org/10.1016/j.bpsgos.2021.09.007
